# CXCL17 is an endogenous inhibitor of CXCR4 via a novel mechanism of action

**DOI:** 10.1101/2021.07.05.451109

**Authors:** Carl W. White, Laura E. Kilpatrick, Natasha Dale, Rekhati S. Abhayawardana, Sebastian Dekkers, Michael J Stocks, Kevin D. G. Pfleger, Stephen J. Hill

## Abstract

CXCL17 is the most recently described chemokine. It is principally expressed by mucosal tissues, where it facilitates chemotaxis of monocytes, dendritic cells, and macrophages and has antimicrobial properties. CXCL17 is also implicated in the pathology of inflammatory disorders and progression of several cancers, as well as being highly upregulated during viral infections of the lung. However, the exact role of CXCL17 in health and disease is largely unknown, mainly due to a lack of known molecular targets mediating CXCL17 functional responses. Using a range of bioluminescence resonance energy transfer (BRET) based assays, here we demonstrate that CXCL17 inhibits CXCR4-mediated signalling and ligand binding. Moreover, CXCL17 interacts with neuropillin-1, a VEGFR2 co-receptor. Additionally, we find CXCL17 only inhibits CXCR4 ligand binding in intact cells and demonstrate that this effect is mimicked by known glycosaminoglycan binders, surfen and protamine sulfate. This indicates that CXCL17 inhibits CXCR4 by a unique mechanism of action that potentially requires the presence of a glycosaminoglycan containing accessory protein. Altogether, our results reveal that CXCL17 is an endogenous inhibitor of CXCR4 and represents an important discovery in our understanding of the (patho) physiological functions of CXCL17 and regulation of CXCR4 signalling.

## Introduction

Chemokines are a large family of small, secreted cytokines that play a central role in the migration of cells. Chemokines are widely expressed throughout the body and facilitate both homeostatic as well as inflammatory immune responses. In addition, chemokines mediate numerous other physiological functions including angiogenesis and organogenesis as well as participate in pathophysiological processes such as cancer progression, autoimmune disorders, and aberrant inflammation^1^. Chemokines induce cellular responses by binding and activating G protein-coupled receptors (GPCRs). This results in the activation of heterotrimeric G proteins followed by downstream signalling, with subsequent β-arrestin recruitment to the chemokine receptor resulting in signal termination via receptor internalisation and desensitization^2^.

CXCL17 is the most recently described chemokine. It is constitutively expressed at high levels in mucosal sites throughout the body and is thought to be involved in the innate immune response via recruitment of monocytes, dendritic cells and macrophages^3,4^. In addition, CXCL17 has antimicrobial properties at high concentrations^5^ and modulates angiogenesis^4,6^. CXCL17 has also been implicated in several pathologies including breast^7^ and hepatocellular cancers^8^, as well as inflammatory lung pathologies such as idiopathic pulmonary fibrosis^5^, asthma^9^, influenza^10^ and SARS-CoV-2^11^ infection. Despite mediating a wide range of important functional responses, the receptor(s) for CXCL17 are still unknown. Thus, it is vital to determine the molecular targets of CXCL17 not only to fully understand the normal function of CXCL17 but to allow targeting of its pathophysiological effects.

Previous work implicated GPR35 as the receptor for CXCL17^12^, however subsequent studies^13,14^ have been unable to reproduce this finding. Despite this, G protein-coupled receptors likely play a role in CXCL17-mediated signalling. Indeed, CXCL17-mediated calcium flux^12^, inhibition of cAMP accumulation^15^ and pertussis toxin sensitive signalling^4,16^ have all been observed. Very recently, we have found CXCL17 is able to activate the atypical chemokine receptor ACKR3 as well as interact with glycosaminoglycans (manuscript in preparation). Since ACKR3 does not activate G protein-mediated signalling^17^, it is unlikely to be the only receptor for CXCL17. Notably ACKR3 interacts closely with the chemokine receptor CXCR4, through formation of ACKR3-CXCR4 heteromers as well as binding and scavenging CXCL12, the cognate ligand for CXCR4^18^.

CXCR4 is a prototypical chemokine receptor that facilitates numerous physiological processes including organogenesis, hematopoiesis, and immune responses via binding of CXCL12. Moreover, CXCR4 signalling is involved in multiple diseases including various cancers, autoimmune disorders, aberrant inflammation and HIV-1 infection^19^. Therefore, the development of inhibitors of CXCR4 is an area of active research^20^. Notably, CXCL12 and CXCR4 are constitutively-expressed at high levels widely throughout the body and several mechanisms exist to regulate CXCR4 signalling. These include: receptor oligomerisation; scavenging of CXCL12 by ACKR3 to regulate ligand localization and abundance; post-translational modifications; and proteolytic degradation of CXCL12^19^. Moreover, positively charged domains within CXCL12 mediate interactions with glycosaminoglycans that facilitate CXCL12 oligomerisation, prevent proteolytic degradation and help establish chemotactic gradients^21^. Finally, several endogenous inhibitors of CXCR4 have now been reported including CXCL14^22^, the endogenous cationic antimicrobial peptide cathelicidin LL37^23^, as well as EPI-X4, an endogenous peptide fragment of serum albumin^24^.

In this study, across a panel of chemokine receptors we find no agonist-like responses mediated by CXCL17. Instead we show that CXCL17 inhibits CXCR4-mediated signalling as well as ligand binding in intact live cells. CXCL17 is therefore an endogenous inhibitor of CXCR4 and we discuss the potential physiological implications of this interaction. Notably, CXCL17 has no effect on binding of CXCL12 or a small molecule antagonist to CXCR4 in dissociated membrane preparations. This suggests that CXCL17 inhibits CXCR4 by a unique mechanism of action which we propose involves an accessory protein containing glycosaminoglycans.

## Results

### GPR35 is not a direct target of CXCL17

In initial confirmatory experiments we sought to rule out GPR35 as a target for CXCL17 using bioluminescence resonance energy transfer (BRET) assays. In HEK293 cells transiently-transfected with GPR35/NLuc and β-arrestin2/Venus (Supplementary Figure 1a) the GPR35 agonist Zaprinast (100 μM) but not CXCL17 (100 nM) resulted in an increase in BRET indicative of recruitment of β-arrestin2/Venus to GPR35/NLuc confirming similar experiments published previously^13,14^. Similarly, in HEK293 cells transiently transfected with GPR35, Gα_i1_/NLuc and Venus/G_γ2_ (Supplementary Figure 1b), no change in the BRET ratio was observed with CXCL17 (100 nM) whereas Zaprinast (100 μM) decreased the BRET ratio in a GPR35 specific and concentration dependent manner (Supplementary Figure 2, pEC_50_ = 4.76 ± 0.82). Next, we examined if activation of GPR35 by CXCL17 required the formation of an oligomeric complex with a chemokine receptor. Here HEK293 cells were transiently-transfected to co-express GPR35, Gα_i1_/Nluc, Venus/G_γ2_ and a chemokine receptor (Supplementary Figure 1c). In each assay configuration the endogenous chemokine ligand (30 nM) resulted in a conformational change of the G protein complex (except for CCR9 for which no response was obtained with the positive control ligand) however CXCL17 (30 nM, sufficient to induce signalling^6^) produced no comparable or observable change in the BRET ratio. Taken together these data confirm GPR35 is not a direct target of CXCL17.

### CXCL17 inhibits CXCR4 signalling

Targets of CXCL17 are likely to be Gα_i_ coupled G protein-coupled receptors due to reports of pertussis toxin sensitive signalling^4,16^. In HEK293 cells transfected with Gα_i1_/NLuc, Venus/Gγ2 and an individual chemokine receptor CCR1-10 or CXCR1-3, 5 or 6 (without co-expression of GPR35), endogenous chemokine ligands resulted in a decrease in the BRET ratio suggestive of receptor activation, however, no such change was observed using a high concentration of CXCL17 (300 nM) either in the absence or presence of endogenous chemokine ligand (Figure 1a, n=4). Surprisingly, while CXCL17 did not induce an agonist-like response, we found that in HEK293 cells co-expressing CXCR4, Gα_i1_/Nluc and Venus/G_γ2_, CXCL17 (300 nM) resulted in an increase in the BRET ratio as well as inhibited responses mediated by CXCL12 (1 nM, Figure 1b) in a concentration-dependent manner (Figure 1c; CXCL12 0.1 nM, pIC_50_ = 6.93 ± 0.28, n=4) albeit with a lower potency than the small molecule CXCR4 antagonist AMD3100 (pIC_50_ = 7.52 ± 0.03, n=4). CXCL17 had similar inhibitory effects in Cos-7 cells transiently transfected with CXCR4, Gα_i1_/NLuc and Venus/G_γ2_, indicating these effects where not cell type dependent (Supplementary Figure 3). To investigate the effect of CXCL17 on downstream CXCR4 signalling we stably-transfected SNAP/CXCR4 into HEK293G cells that express the GloSensor cAMP biosensor. In this system CXCL17 and the small molecule CXCR4 antagonist AMD3100 inhibited the CXCL12-mediated decrease in forskolin-induced cAMP accumulation (Figure 1d, p<0.01, n=4, one way ANOVA with Dunnett’s multiple comparisons test). Interestingly, in the Gα_i1_/NLuc and Venus/G_γ2_ activation assay using a constitutively-active CXCR4 mutant (N119S), we observed an increase in the magnitude of inhibition (compared to at wildtype CXCR4) mediated by a submaximal concentration of CXCL17 (100 nM, p<0.05, WT CXCR4 versus N119S CXCR4, n=3, two-tailed unpaired t-test), whereas AMD3100 (1 μM) appeared to mediate a small decrease in BRET indicative of receptor activation and is in keeping with reports of AMD3100 being a partial agonist for constitutively active CXCR4^25^ (Figure 1e). Finally, N-terminal truncation of chemokines can drastically modulate function, indeed an increase in CXCL17 potency has been observed following N-terminal truncation^6^. 24Leu-CXCL17 has been used in most reports and is the canonical version of CXCL17 we have used here. However, in a direct comparison we observed no difference in the ability of 24LeuCXCL17 or a commercially available truncated version 22Ser-CXCL7 to inhibit constitutive CXCR4 mediated changes in Gα_i1_/NLuc-Venus/G_γ2_ conformation in HEK293 cells (Figure 1f, p>0.05, n=3). Similarly, 22Ser-CXCL17 (300 nM) still inhibited the decrease in BRET mediated by CXCL12 (1 nM, Figure 1f, p<0.05, n=3). Altogether these results suggest CXCL17 selectively inhibits CXCR4 signalling and has a different mechanism of action compared to small molecule antagonists.

**Figure 1:**
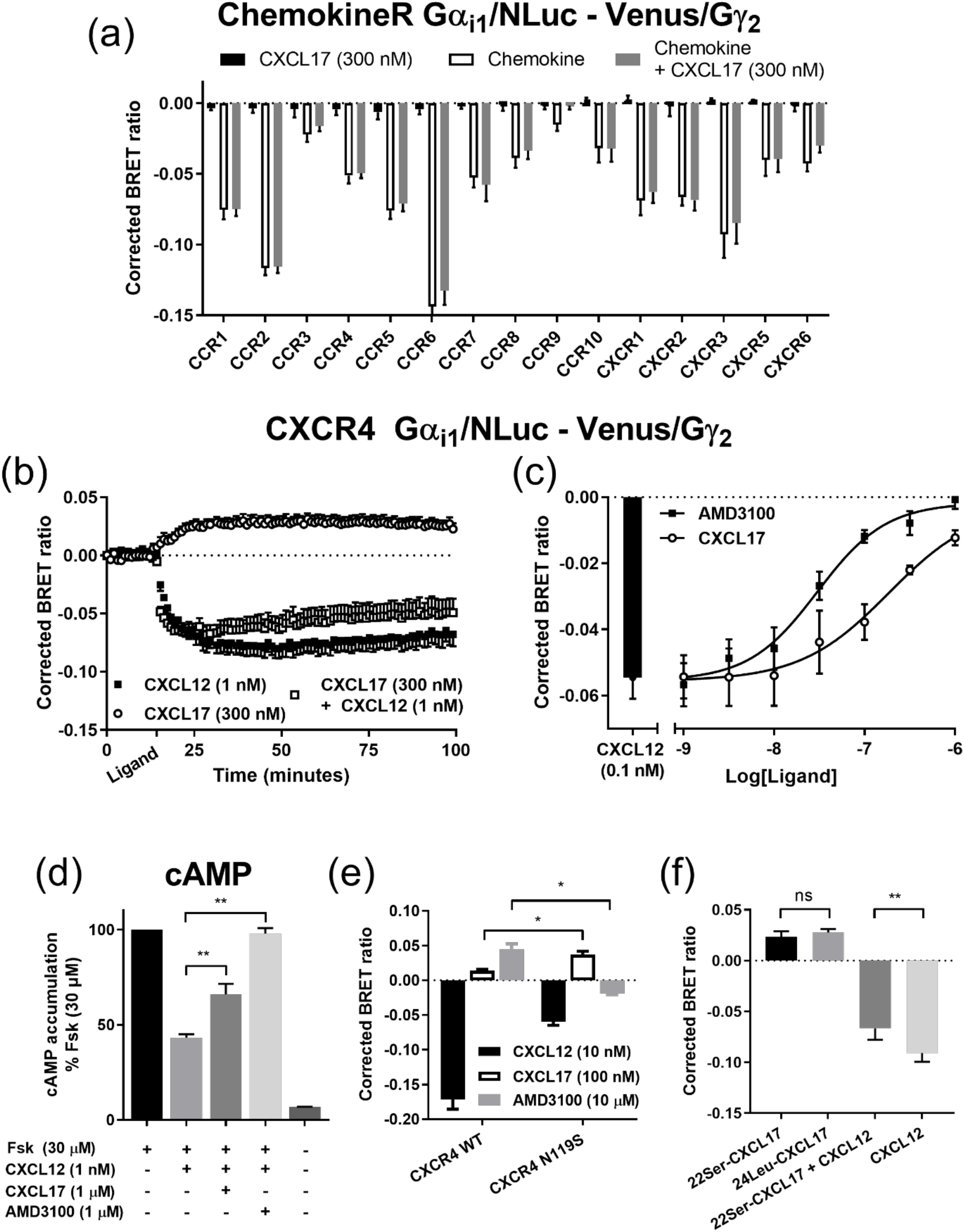
CXCL17 inhibits constitutive, and ligand-induced CXCR4 signalling. **(a)** HEK293 cells transiently transfected with CCR1-10 or CXCR1-3,5 or 6, Gα_i1_/Nluc and Venus/G_γ2_ were stimulated with CXCL17 (300 nM, black bars) or submaximal concentrations of chemokine CCL3 (0.3 nM), CCL2 (0.3 nM), CCL13 (1 nM), CCL22 (0.3 nM), CCL4 (30 nM), CCL20 (3 nM), CCL19 (1 nM), CCL1 (10 nM), CCL25 (10 nM), CCL27 (10 nM), CXCL8 (0.3 nM), CXCL8 (1 nM), CXCL11 (1 nM), CXCL13 (3 nM) and CXCL16 (1 nM) for CCR1-10 and CXCR1-3,5 and 6 respectively or CXCL17 (300 nM) and the corresponding chemokine ligand (grey bars). **(b)** Kinetic analysis of changes in BRET following application of CXCL12 (1 nM, black squares), CXCL17 (300 nM, white circles) or both CXCL12 and CXCL17 (1 nM and 300 nM respectively, white squares) in HEK293 cells transiently-transfected with CXCR4, Gα_i1_/Nluc and Venus/G_γ2_. **(c)** Inhibition of the change in BRET mediated by CXCL12 (0.1 nM) in HEK293 cells transiently transfected with CXCR4, Gα_i1_/Nluc and Venus/G_γ2_ by increasing concentrations (1 nM – 1 μM) of AMD3100 (black squares) or CXCL17 (white circles). **(d)** HEK293G cells stably-expressing the cAMP GloSensor transgene and SNAP/CXCR4 were stimulated with buffer, or forksolin (30 μM), or forskolin (30 μM) and CXCL12 (1 nM), in the absence or presence of CXCL17 (1 μM) or AMD3100 (1 μM). **(e)** HEK293 cells transiently-transfected with Gα_i1_/Nluc, Venus/G_γ2_ and wildtype (WT) CXCR4 or the CXCR4 N119S constitutively-active mutant were stimulated with CXCL12 (10 nM, black bars), CXCL17 (100 nM, white bars) or AMD3100 (10 μM, grey bars). **(f)** In parallel, HEK293 cells transiently-transfected with CXCR4, Gα_i1_/Nluc and Venus/G_γ2_ were stimulated with canonical CXCL17 (24Leu-CXCL17, 300 nM), a truncated version of CXCL17 (22Ser-CXCL17, 300 nM), CXCL12 (1 nM), or 22Ser-CXCL17 (300 nM) and CXCL12 (1 nM). For **b,** ligand was added following establishment of basal BRET and is indicated on the x-axis as *ligand*. Corrected BRET was calculated as described in *Methods*. Points or bars represent mean ± s.e.m. of three (**e** and **f**) or four (**ad**) individual experiments performed in duplicate or triplicate. **(d)** bars represent mean % ± s.e.m. of the forskolin-mediated response of each individual experiment. *, p<0.05 and **, p<0.01, indicates a significant difference between the paired groups. Statistical analysis by **(d)** one-way ANOVA with a Dunnett’s multiple comparisons test or **(e-f)** paired t-test.

### CXCL17 inhibits CXCR4-β-arrestin2 interactions

To confirm the inhibitory effects of CXCL17 we next investigated its effect on recruitment of β-arrestin-2 to the CXCR4. In HEK293 cells transiently transfected with CXCR4/Rluc8 and β-arrestin2/Venus, application of CXCL17 (100 nM, Figure 2a, n=3) resulted in a decrease in the basal BRET ratio as well as inhibition of CXCL12 (100 nM) mediated increases in BRET, thus indicating CXCL17 inhibits both constitutive and ligand-induced CXCR4 activity. The decrease in the basal BRET ratio following the application of CXCL17 was concentration-dependent (Figure 2b, pIC_50_ = 7.24 ± 0.08, n= 3) and in HEK293 cells transfected with CXCR4/NLuc and β-arrestin2/Venus (Figure 2c) both CXCL17 (pIC_50_ = 6.85 ± 0.30, n=6) and AMD3100 (pIC_50_= 7.79 ± 0.14, n=6) inhibited the increase in BRET mediated by CXCL12 (10 nM) in a concentration-dependent manner. To further test the specificity of CXCL17 for CXCR4, we used HEK293 cells transfected with CXCR1/Rluc8, CCR5/Rluc8 or β2-adrenoceptor/Rluc8 (β2-AR) and β-arrestin2/Venus. Each configuration showed an increase in BRET following application of a prototypical agonist but no modulation of the BRET ratio following application of CXCL17 (300 nM, Figure 2d, n=3).

**Figure 2:**
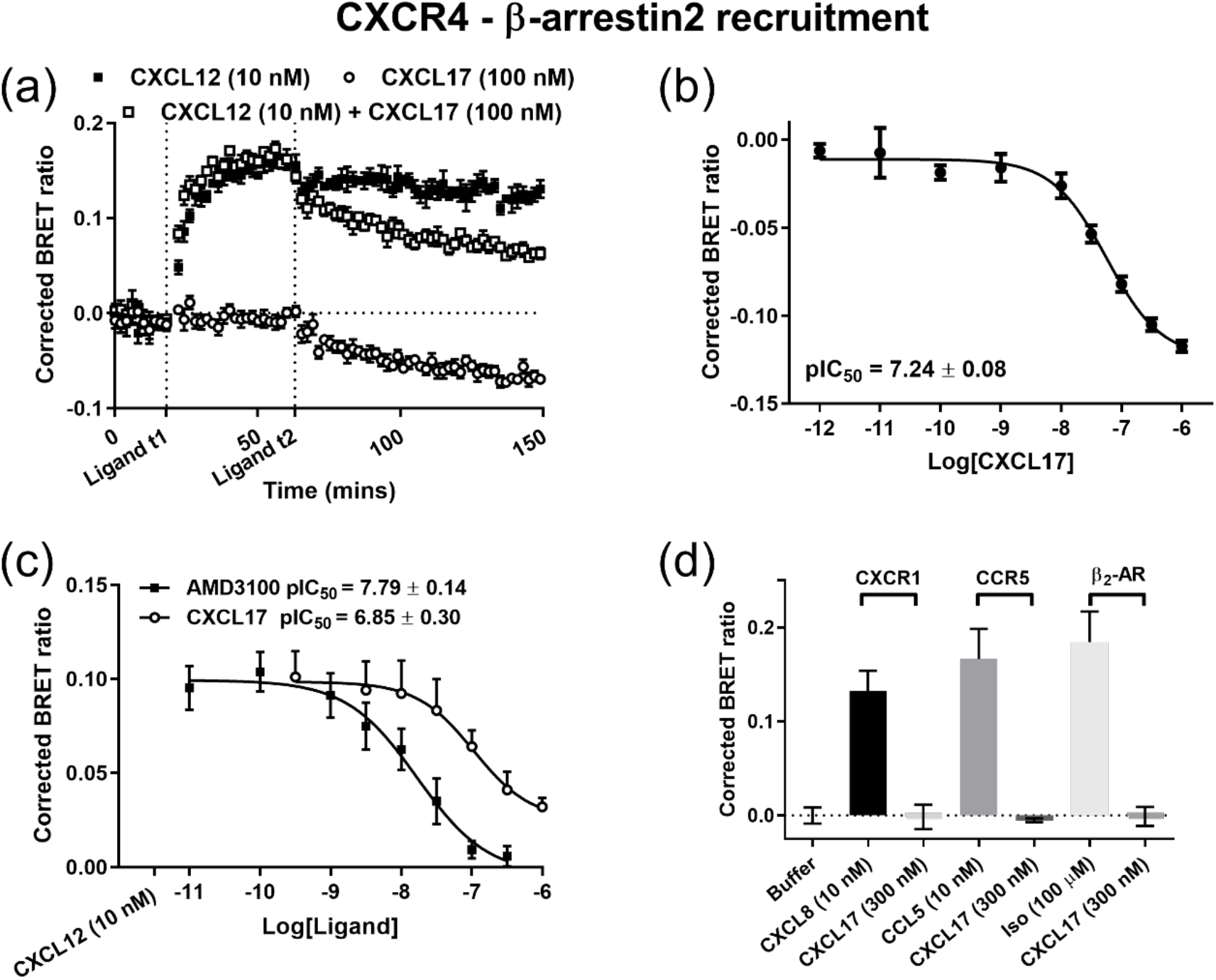
CXCL17 inhibits constitutive, and ligand-induced CXCR4-β-arrestin2 interactions. **(a)** HEK293 cells transiently-transfected with CXCR4/Rluc8 and β-arrestin2/Venus were stimulated at time *ligand t1* with HBSS (white circles) or CXCL12 (100 nM, black and white squares) then at time *ligand t2* stimulated again with HBSS (black squares) or CXCL17 (300 nM, white squares and circles) and change in BRET observed. **(b**) Change in BRET following application of increasing concentrations of CXCL17 (1 pM – 1 μM) to HEK293 cells transiently-transfected with CXCR4/Rluc8 and β-arrestin2/Venus. **(c)** HEK293 cells transiently-transfected with CXCR4/NLuc and β-arrestin2/Venus were stimulated with CXCL12 (10 nM) in the absence (black bar) or presence of AMD3100 (black squares) or CXCL17 (white circles). **(d)** HEK293 cells transiently-transfected with CXCR1/Rluc8, CCR5/Rluc8 or β_2_-adrenoceptor/Rluc8 and β-arrestin2/Venus were stimulated with CXCL17 (300 nM) or either CXCL8 (10 nM), CCL5 (10 nM) or isoprenaline (100 μM) respectively. Corrected BRET was calculated as described in *Methods*. Points or bars represent mean ± s.e.m. of three (**a, b and d**) or six (**c**) individual experiments performed in duplicate or triplicate and points and bars in (**b-d**) represent the maximum change in BRET from a kinetic experiment.

### CXCL17 inhibits ligand binding to NLuc/CXCR4 in a context dependent manner

Initial observations suggested that CXCL17 had a unique mechanism of action/binding compared to known antagonists. To further examine binding of CXCL17 to CXCR4 we used a NanoBRET ligand binding approach. In live HEK293 cells stably-expressing exogenous CXCR4 tagged on the N-terminus with NanoLuc (NLuc/CXCR4), both AMD3100 and CXCL17 competed with CXCL12-AF647 (12.5 nM, Figure 3a, pKi = 7.99 ± 0.16 and pKi = 6.25 ± 0.19 respectively, n=4) for NLuc/CXCR4 binding in a concentration-dependent manner. Recent studies have demonstrated the oligomeric state of CXCR4 is expression dependent and that oligomer formation and signalling can be disrupted by ligands such as IT1t that bind allosterically^26,27^. However, when binding experiments were performed in live HEK293 or HeLa cells CRISPR/Cas9 genome-edited to express NLuc/CXCR4 under endogenous promotion and therefore with native levels of CXCR4 expression (Supplementary Figure 4), AMD3100 (pKi = 8.01 ± 0.13 and pKi = 8.30 ± 0.08 respectively, n=4) and CXCL17 (pKi = 6.14 ± 0.30 and pKi = 6.15 ± 0.27 respectively, n=4) inhibited CXCL12-AF647 (12.5 nM) binding. These results rule out that this effect was specific to over-expressed CXCR4 receptors, potentially existing predominantly as CXCR4 oligomers, and further demonstrating inhibition was independent of cell type. Indeed, supporting a unique binding mode, CXCL17 did not compete with CXCL12-AF647 binding to NLuc/CXCR4 in membrane preparations from HEK293 cells stably-overexpressing the receptor (Figure 3b, n=4) whereas AMD3100 inhibited CXCL12-AF647 (12.5 nM) binding with an affinity similar to that observed in whole live cells (Figure 3b, pKi = 7.84 ± 0.06, n=4).

**Figure 3:**
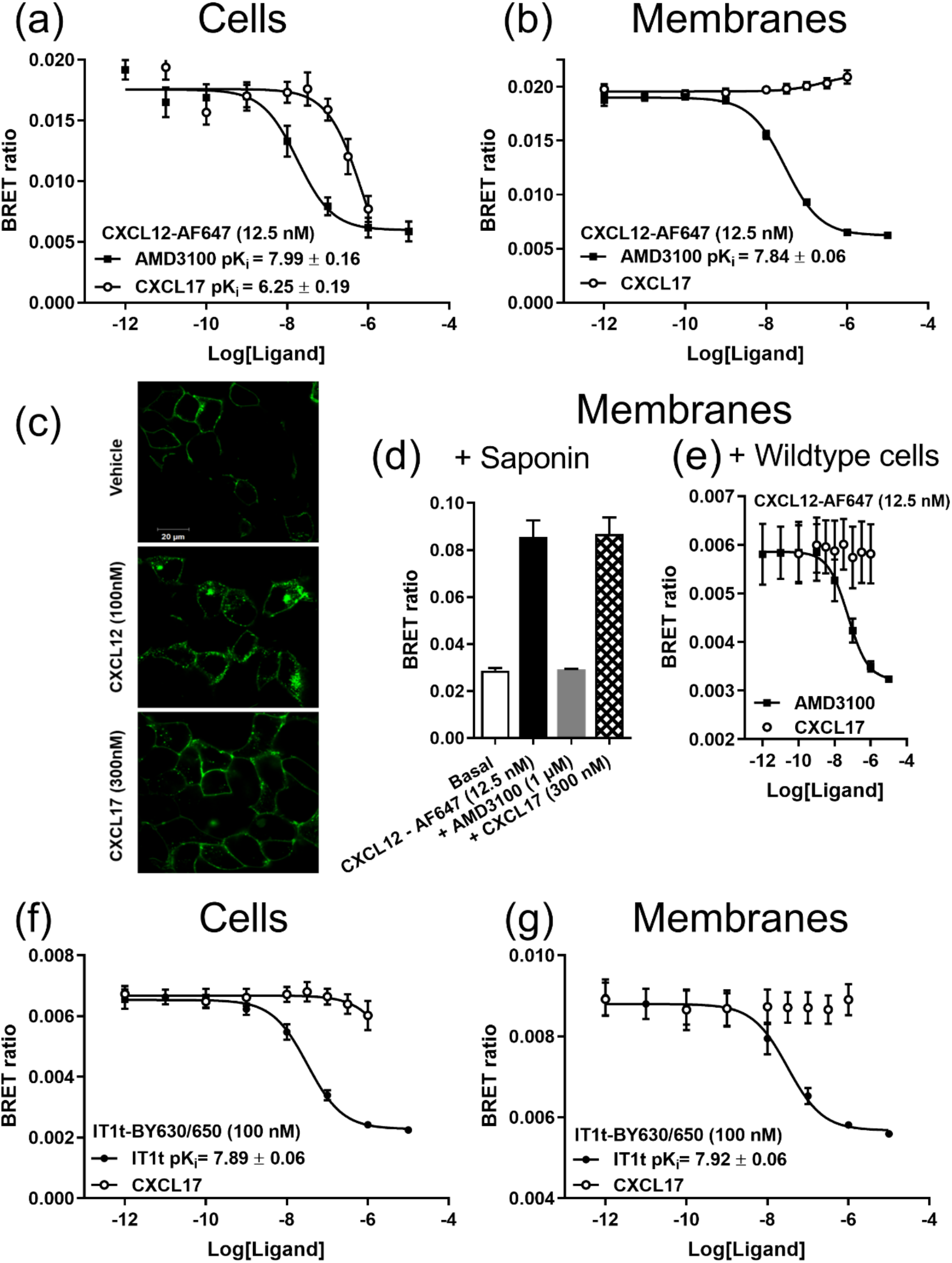
CXCL17 binding to CXCR4 is context dependent. Live **(a)** HEK293 cells or **(b)** membrane preparations stably-expressing NLuc/CXCR4 were incubated for 1 hr at 37 °C with CXCL12-AF647 (12.5 nM) and increasing concentrations of AMD3100 (10 pM – 10 μM, black squares) or CXCL17 (100 pM – 1 μM, white circles). **(c)** Confocal imaging (Zeiss LSM 710) of HEK293 cells stably-expressing SNAP/CXCR4 under unstimulated conditions (vehicle) or after treatment with 100 nM CXCL12 or 300 nM CXCL17 (1 h at 37°C). Data are representative of three individual experiments. Scale bar represents 20 μm. **(d)** Membrane preparations stably-expressing NLuc/CXCR4 were permeabilised with saponin (0.25 mg/ml) and incubated for 1 hr at 37 °C with CXCL12-AF647 (12.5 nM) in the absence (black bar) or presence of AMD3100 (1 μM, grey bar) or CXCL17 (300 nM, hatched bar) white bar (HBSS) is vehicle control. **(e)** Membrane preparations stably expressing NLuc/CXCR4 were co-incubated with wildtype (untransfected) HEK293 cells and incubated for 1 hr at 37 °C with CXCL12-AF647 (12.5 nM) and increasing concentrations of AMD3100 (10 pM – 10 μM, black squares) or CXCL17 (100 pM – 1 μM, white circles). **(f)** HEK293 cells or **(g)** membrane preparations stably-expressing NLuc/CXCR4 were incubated for 1 hr at 37 °C with IT1t-BY630/650 (100 nM) and increasing concentrations of AMD3100 (10 pM – 10 μM, black squares) or CXCL17 (100 pM – 1 μM, white circles). Points or bars represent mean ± s.e.m. of four (**a, b, d, e and f)**) or five **(g)** individual experiments performed in duplicate.

Differences in binding between whole cells and membranes could plausibly be due to internalisation and thus compartmentalisation of the receptor. In live HEK293 cells stably-expressing SNAP/CXCR4 labelled with cell-impermeant AlexaFluor488, application of CXCL12 (100 nM) but not CXCL17 (300 μM) for 30 minutes at 37°C resulted in extensive receptor internalisation (Figure 3c). Furthermore, using a Nanoluciferase complementation assay where live cells expressing CXCR4 tagged on the N-terminus with HiBiT (HiBiT/CXCR4) are complemented with a cell impermeant fragment of NanoLuc (LgBiT), CXCL17 resulted in an increase in luminescence at high concentrations (>100 nM) indicating accumulation of HiBiT/CXCR4 at the plasma membrane not internalisation (Supplementary Figure 5a). By taking advantage of the cell impermeant nature of the purified fragment of NanoLuc we were able to confine ligand binding to the plasma membrane and confirmed that CXCL17 addition still resulted in the displacement of CXCL12-AF647 (12.5 nM) with similar affinity (Supplementary Figure 5b, pKi = 6.60 ± 0.1, n=4) in live cells even when agonist-induced internalisation and compartmentalisation was removed. Similarly, in membrane preparations permeabilised with 0.25 mg/ml saponin (Figure 3d), AMD3100 (1 μM) but not CXCL17 (300 nM) displaced CXCL12-AF647 (12.5 nM), ruling out that membrane vesicle formation was resulting in ligand-receptor compartmentalisation. Next, to investigate if more active forms of CXCL17 were being produced due to cleavage by cellular proteases in live cells, membrane preparations expressing NLuc/CXCR4 were co-incubated with wildtype HEK293 cells and binding investigated. However, no inhibition of CXCL12-AF647 (12.5 nM) binding by CXCL17 (300 nM) was observed (Figure 3e).

Finally, on close inspection of the competition ligand binding curves (Figures 3a-b) we noted a steep slope >1 suggestive of non-competitive and/or allosteric interactions and therefore hypothesised CXCL17-mediated ligand displacement would be probe-dependent. To this end, the CXCR4 antagonist IT1t but not CXCL17 competed with binding of IT1t-BY630/650 (a fluorescent derivative of IT1t, pK_d_ = 7.25 ± 0.14, n=5, Supplementary Figure 6) to NLuc/CXCR4 expressed in live HEK293 cells or to NLuc/CXCR4 in membrane preparations (Figure 3f-g, pKi = 7.89 ± 0.06 and pKi = 7.92 ± 0.06 for IT1t in cells and membranes respectively, n=4-5). Since the binding site of IT1t only partially overlaps with CXCL12^28^, these results further indicate that CXCL17 binds by a unique mechanism/site compared to the endogenous agonist CXCL12 and other known CXCR4 antagonists.

### CXCL17 prevents endogenous CXCL12 binding to CXCR4

In our initial experiments we also noted that CXCL17 inhibits basal/constitutive CXCR4 signalling, suggesting inverse agonist activity. However, CXCL12 is endogenously expressed in HEK293 cells at levels sufficient to initiate signalling^29^. To investigate if CXCL17 was disrupting signalling mediated by endogenous CXCL12, we took advantage of HEK293 cells engineered using CRISPR/Cas9 genomeediting to express CXCL12 tagged with a small 11 amino acid self-complementing fragment of NLuc (HiBiT). When the cells were transfected with exogenous SNAP/CXCR4, AMD3100 and CXCL17 both produced a concentration-dependent decrease in BRET (Figure 4, pIC_50_ = 7.10 ± 0.08 and 6.27 ± 0.16 respectively, n=6), generated between CXCL12-HiBiT complemented with LgBiT and SNAP/CXCR4 labelled with cell-impermeant AlexaFluor488. These data indicate that inhibition of constitutive CXCR4 signalling is likely due to blocking endogenous CXCL12 binding to CXCR4.

**Figure 4:**
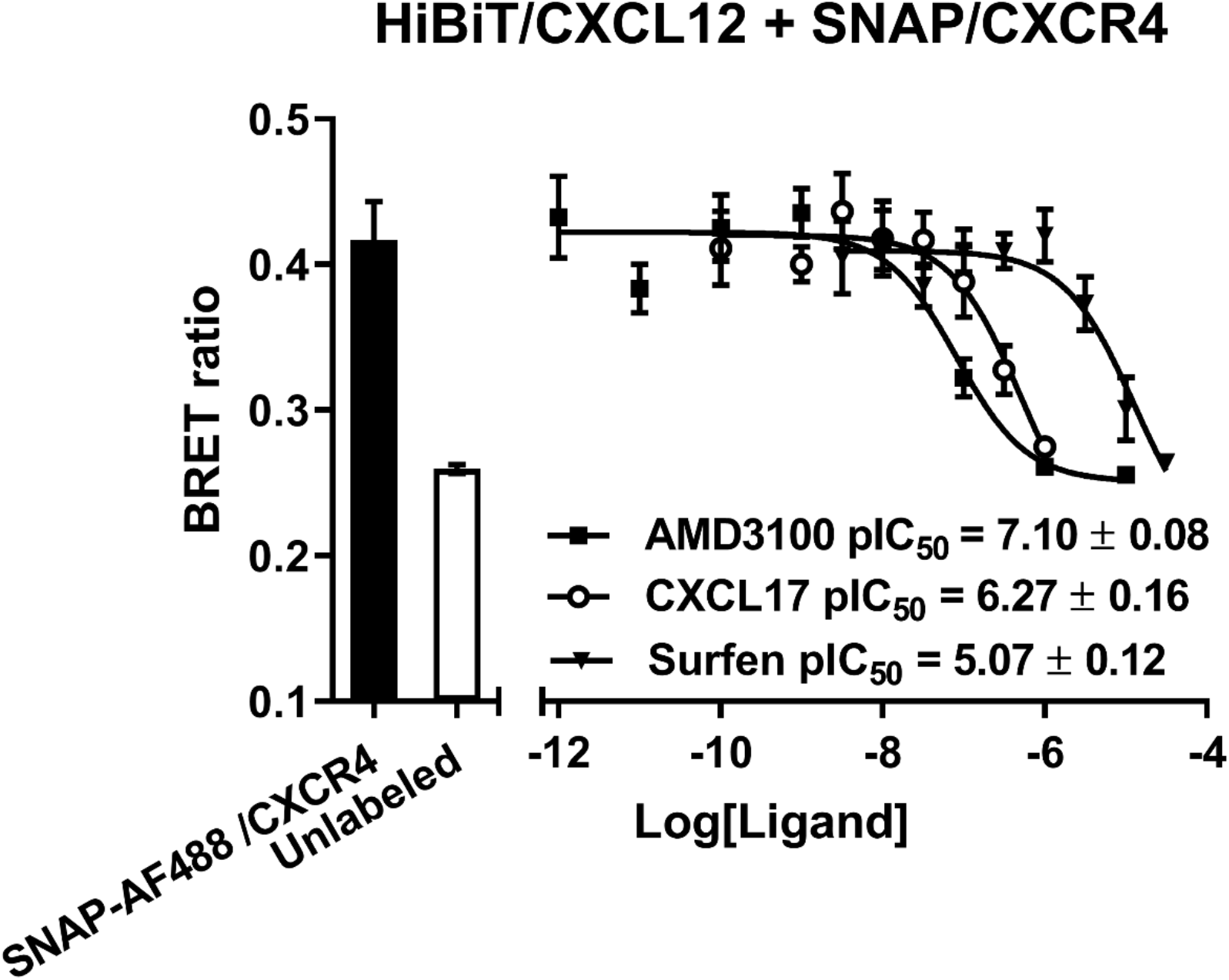
Displacement of the binding of secreted genome-edited CXCL12-HiBiT to SNAP/CXCR4 by CXCL17. HEK293 cells expressing genome-edited CXCL12-HiBiT, were transiently transfected with SNAP/CXCR4 were incubated for 1 hr at 37 °C in the absence or presence of increasing concentrations of AMD3100 (black squares, 100 pM – 10 μM), CXCL17 (white circles, 100 pM – 1 μM) or Surfen (black triangles, 30 nM – 30 μM). CXCL12-HiBiT was complemented with purified LgBiT (30 nM) and SNAP/CXCR4 labelled with cell impermeant AlexaFluor488 prior to measurement of BRET. Bars represent basal BRET in the absence of added AlexaFluor488 label. Bars and points represent mean ± s.e.m. of six individual experiments performed in duplicate.

### Evidence for the involvement of glycosaminoglycans in CXCL17 inhibition of CXCR4

Chemokines in addition to membrane receptors bind GAGs via basic domains (putatively BBxB) that result in chemokine accumulation at the cell surface and facilitates oligomerisation as well as prevents proteolytic degradation. CXCL17 has an overall positive net charge of +18 at pH 7.4 and contains multiple highly conserved (Supplementary Figure 7) putative GAG domains (Figure 5a) indicating an ability to bind GAGs potentially with high affinity. As generation of membrane preparations undoubtedly disrupts the extracellular matrix, we therefore hypothesised that CXCL17 required the presence of (unperturbed) membrane-bound glycosaminoglycans to inhibit CXCR4. To test this, we determined whether the effect of CXCL17 could be mimicked by known GAG binders. Both the small molecule GAG inhibitor surfen and the heparin antidote protamine sulfate (previously shown to inhibit CXCR4 signalling^30^) inhibited CXCL12-AF647 binding to NLuc/CXCR4 in cells (Figure 5b-c, (pK_i_ = 4.85 ± 0.61 and pK, = 6.35 ± 0.13, surfen and protamine sulfate respectively) but not in membranes (Figure 5d-e). Further confirming that GAG binders can modulate CXCR4, surfen displaced CXCL12-HiBiT binding to SNAP/CXCR4 in live HEK293 cells in a concentration-dependent manner (Figure 4, pIC_50_ = 5.07 ± 0.12). Consistent with the effects mediated by CXCL17, the slope of the concentration response curves of both GAG inhibitors was greater than 1 indicating an allosteric and/or noncompetitive interaction. Notably, and as seen with CXCL17, surfen did not compete with IT1t-BY630/650 binding to NLuc/CXCR4 in live cells (Supplementary Figure 8a). Next, we used exogenous soluble heparan sulphate to pre-occupy the GAG binding sites of CXCL17 and therefore reduce its capacity to bind to endogenous GAG in the extracellular matrix. We observed that while heparan sulfate (30 μg/ml) treatment had no effect on the binding of CXCL12-AF647 (12.5 nM) to NLuc/CXCR4 in live HEK293 cells (Supplementary Figure 8b), preincubation of CXCL17 (300 nM) with heparan sulfate (30 μg/ml) reduced the ability of CXCL17 to cause displacement of CXCL12-AF647 binding to NLuc/CXCR4 (p<0.01, n=5) whereas heparan sulfate had no effect on AMD3100 (1 μM). Finally, we investigated if inhibition of CXCR4 ligand binding was a general property of high affinity GAG binders. Using a high concentration of CXCL4 (1 μM, a.k.a. platelet factor 4, GAG affinity ~ 30 pM^31^) we did not observe inhibition of CXCL12-AF647 (12.5 nM) binding to NLuc/CXCR4 in live HEK293 cells (Supplementary Figure 8c). These results suggest GAGs are involved in the CXCL17 mediated inhibition and binding of CXCR4, but that not all known GAG binders can inhibit binding of CXCL12 to CXCR4.

**Figure 5:**
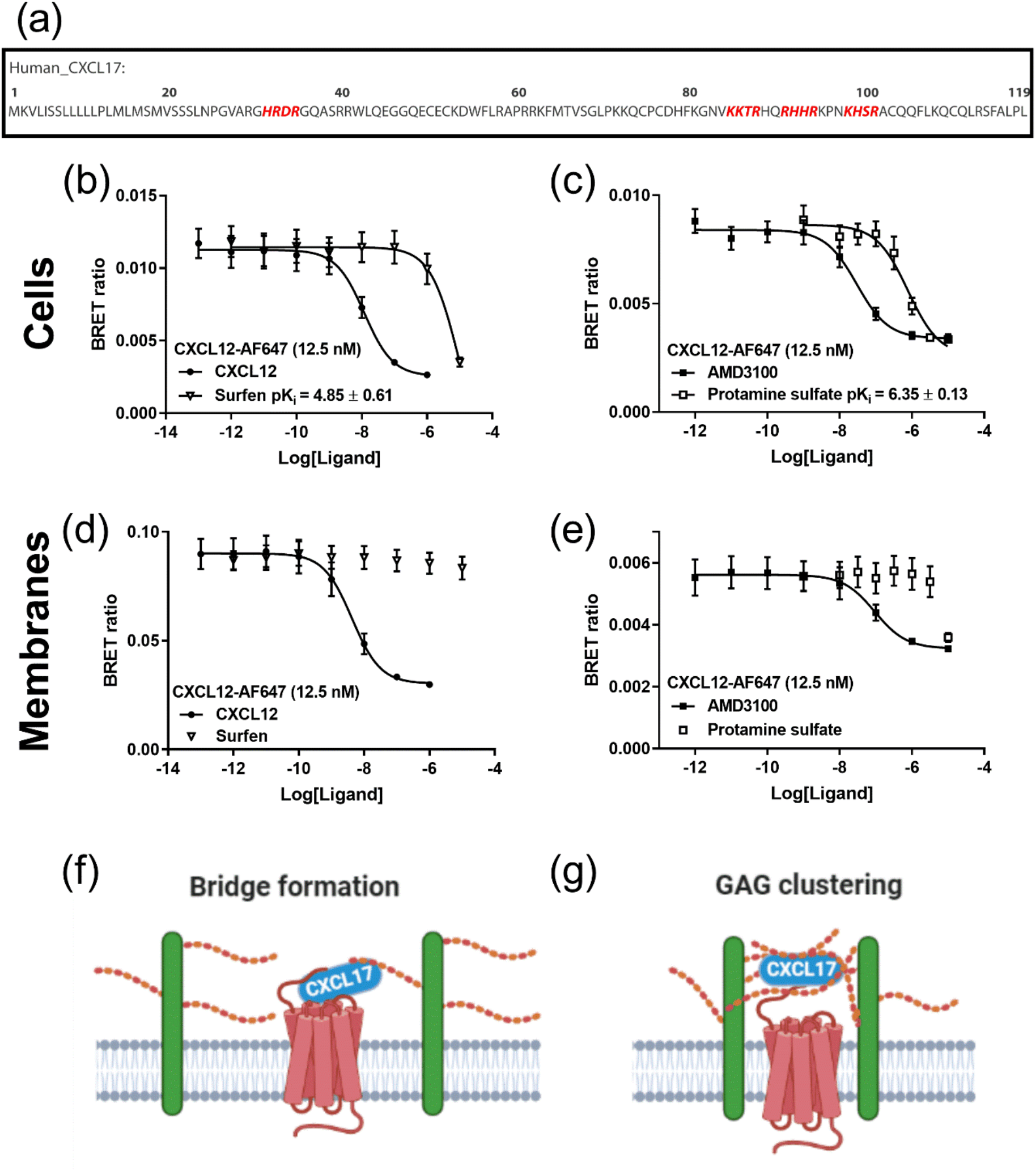
Binders of glycosaminoglycans mimic the effect of CXCL17. **(a)** Sequence of human CXCL17 with putative GAG binding domains highlighted in red. **(b-e) effect of GAG binders on ligand binding to Nluc/CXCR4.** Live HEK293 cells **(b and c)** or membrane preparations **(d and e)** expressing Nluc/CXCR4, were incubated for 1 hr at 37 °C with CXCL12-AF647 (12.5 nM, **b-e**) and increasing concentrations of surfen (white triangles, 10 pM – 10 μM, **b and d**) or protamine sulfate (white squares, 1 nM – 10 μM, **c and e**), CXCL12 (black circles, 0.1 pM-1 μM, **b and d**) or AMD3100 (black squares, 1pM – 10 μM, **c and e**). Points represent mean ± s.e.m. of four (**b and d**) or five **(c and e)** individual experiments performed in duplicate. **(f-g) Proposed mechanisms of CXCL17-mediated inhibition of CXCR4. (f)** Inhibition of CXCR4 by CXCL17 ‘bridge’ formation between a closely colocalised GAG-containing proteoglycan and CXCR4. **(g)** CXCL17-promoted clustering of GAG containing proteoglycans around CXCR4 prevents binding of CXCL12 but allows binding of small molecule inhibitors.

While GAGs are important regulators of chemokine function *in vivo*, it is hypothesised that they are not strictly required for receptor binding. Indeed, dual GAG-receptor binding appears sterically unlikely for most chemokines^32^. In contrast, GAG binding motifs are required for ligand binding to some receptor tyrosine kinases^33^. Indeed, the vascular endothelial growth factor receptor 2 (VEGFR2) and its co-receptor neuropilin-1 NRP1 both bind full length VEGF-A, however alternatively spliced isoforms of VEGF that lack a GAG binding motif do not bind to NRP1^34,35^. Using this model, we hypothesised that CXCL17 would only inhibit ligand-receptor binding where ligand-GAG interactions are a stringent requirement. Using NanoBRET ligand binding in HEK293 cells transiently-transfected with NLuc/VEGFR2 (Figure 6a) binding of a single site fluorescently-labeled version of VEGF (VEGF165a-TMR; 3 nM^35,36^) that contains the putative GAG binding motif was displaced by unlabelled VEGF165a (30 nM) but not surfen (30 μM), CXCL17 (300 nM) or CXCL12 (100 nM). In contrast in HEK293 cells transiently-transfected with NLuc/NRP1 (Figure 6b), VEGF165a-TMR (3 nM) binding was displaced by unlabelled VEGF165a (30 nM), surfen (30 μM) and CXCL17 (300 nM) but not CXCL12 (100 nM). Taken together, these results further support the involvement of GAG binding motifs in the binding mechanism of CXCL17 and suggests a direct interaction with VEGF signalling pathways.

**Figure 6:**
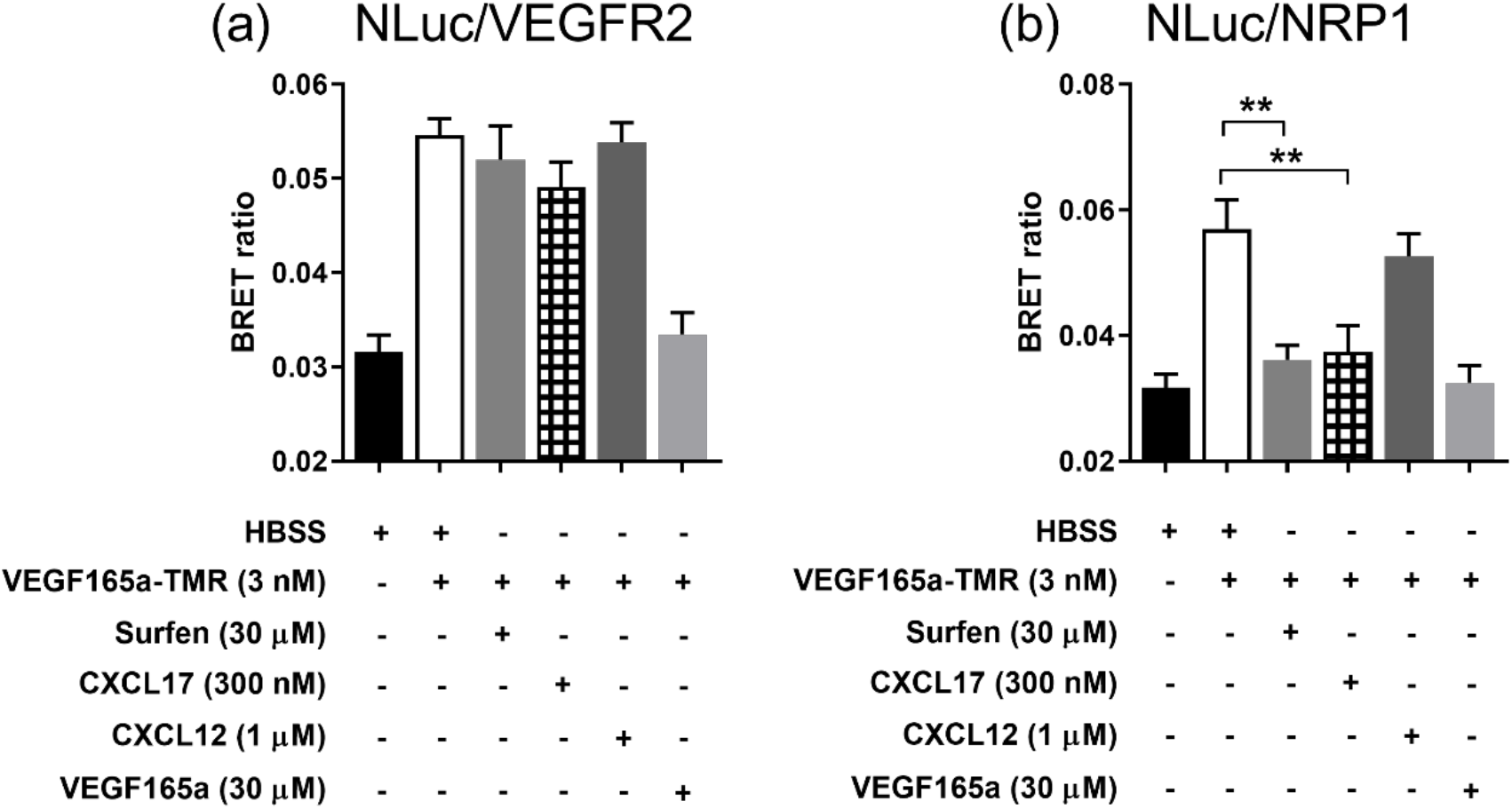
CXCL17 inhibits VEGF165a-TMR binding to NRP1 but not VEGFR2. NanoBRET competition ligand binding in live HEK293 cells expressing **(a)** NLuc/VEGFR2 or **(b) NLuc**/NRP1. Cells were incubated with VEGF165a-TMR (3 nM) in the absence of other ligands (white bar) or in the presence of Surfen (30 μM, grey bar), CXCL17 (300 nM, hatched bar), CXCL12 (1 μM, dark grey bar) or VEGF165a (30 μM, light grey bar). Black bar (HBSS) represents vehicle control. Bars represent mean ± s.e.m. of four individual experiments performed in duplicate. **, p<0.01 calculated by one-way ANOVA with Dunnett’s multiple comparisons test.

## Discussion

CXCL17 was first described in the literature as having chemotactic properties for dendritic cells and monocytes^4^, as well as being correlated with VEGF expression^37^. Since then, our knowledge of CXCL17 in both health and disease has expanded to include functions as an important innate immune factor at mucosal barriers, regulation of angiogenesis, and involvement in tumorigenesis (for review see^38^). However, our current understanding is limited by a lack of *bonafide* molecular targets via which CXCL17 binds and elicits its function. Previously, GPR35, an orphan G protein-coupled receptor, was reported to be a receptor for CXCL17^12^. However, multiple investigations^13,14^, including those presented here have failed to observe similar CXCL17-GPR35 interactions.

Here we report an inhibitory interaction between CXCL17 and CXCR4 across multiple assays, cell lines and contexts. These results contrast with previous studies that found no effect on CXCR4 signalling or ligand binding by CXCL17^6,12^. However, this is unsurprising given the fact that the focus of the previous investigations was on agonist-induced signalling (unlike the antagonistic responses seen here), and that previous ligand binding studies used membrane preparations which, as demonstrated in the current study, do not permit CXCL17 binding to CXCR4. *In vivo*, endogenous CXCR4 inhibitors may represent an important regulatory mechanism to prevent excessive receptor activation. Notably CXCL17 expression can reach ~ 60 ng/ml *in vitro* while serum CXCL17 concentrations can reach ~ 5 ng/ml during influenza infection^10^. Since CXCL17 expression is highly localised to secretory cells in mucosal tissue ^39^ it is plausible that CXCL17 expression reaches concentrations sufficient to inhibit CXCR4.

When compared to AMD3100, which has been reported to be a weak partial CXCR4 agonist^25^, and the inverse agonist IT1t^40^, CXCL17 has a unique mechanism by which inhibition of CXCR4 is achieved. Here the most striking evidence for this is the inability of CXCL17 to inhibit CXCL12-AF647 binding to CXCR4 in membrane preparations. We initially considered that a lack of CXCL17-mediated effects in membranes preparations was due to partitioning or internalisation of CXCR4 in cells. However, limiting ligand binding to the plasma membrane using HiBiT-tagged CXCR4 or cell permeabilization did not support such a mechanism. We also found that CXCL17 did not displace a fluorescently-labeled derivative of IT1t from Nluc/CXCR4 in whole live cells or membrane preparations which could suggest that the formation of CXCL12-AF647-CXCL17 interactions (heteromerization) contributed to CXCL17 inhibition of CXCR4. However, such an interaction would also be expected to occur in membrane preparations when co-incubated with live cells.

In our efforts to delineate the mechanism of action of CXCL17 we noted a steep slope >1 of CXCL17-mediated displacement of CXCL12-AF647 in the NanoBRET competition ligand binding curves which is indicative of an allosteric and/or non-competitive interaction. Furthermore, since CXCL17 had no effect in membrane preparations this suggested the involvement of a site that was lost once intact cells were disrupted and pointed to the involvement of an accessory protein. Here we present several lines of evidence that this potential accessory protein contains glycosaminoglycans: 1) CXCL17 contains multiple highly conserved putative GAG binding domains and preparation of purified cell membranes certainly disrupts the native structure, function and/or potentially presence of GAGs^41^. 2) In support of this, the effect of CXCL17 was mimicked by known GAG binders, surfen and protamine sulphate. Further evidence from the present study for the involvement of GAGs are: a) heparan sulfate reduced CXCL17-mediated inhibition of CXCL12-AF647 NLuc/CXCR4 binding presumably via blockade of CXCL17 GAG binding domains and b) CXCL17 inhibited binding of VEGF_165_a-TMR binding to NLuc/NRP1 but not NLuc/VEGFR2. GAG binding domains are strictly required for VEGF_165_a binding to NRP but not for binding to VEGFR2^35^. Notably there does appear to be a level of specificity for CXCL17-mediated inhibition of CXCR4 since the high affinity GAG binder CXCL4^31^ did not inhibit CXCR4, and CXCL17 does not broadly inhibit or activate other chemokine receptors. While yet to be determined, such specificity may be encoded through a dependence on the composition of the GAG side chains (i.e. heparan sulfate versus chondroitin sulfate) for CXCL17 binding, the need for CXCL17 to contain both GAG and CXCR4 binding domains, or inhibition of CXCR4 may depend on interactions with a specific membrane proteoglycan to bring specific GAG binding domains in close proximity with CXCR4. For example, in smooth muscle cells NRP1 is post-translationally modified by N-linked glycosylation and by the addition of *O*-linked chondroitin sulphate to serine 612 (S612) whilst in endothelial cells it is derivatised with heparan sulphate^42^.

A plausible explanation for these results could be that CXCL17 participates in the release of an endogenous factor that subsequently binds and inhibits CXCR4. Indeed, competitive chemokine binding for GAGs is known to induce cooperative signalling due to the ‘release’ of GAG bound chemokines subsequently activating a secondary receptor^43^. However, CXCL17 had no effect on CXCL12-CXCR4 interactions when membrane preparations expressing NLuc/CXCR4 were coincubated with live intact wildtype HEK293 cells, making such an effect unlikely. In contrast, when considering the mechanism of CXCL17 inhibition of CXCR4, these results principally support trans-interactions that are between CXCR4 and a proteoglycan/GAG on the same cell rather than cis interactions between receptors and GAGs on different cells. Thus, the simple presence of GAGs is not sufficient for CXCL17-mediated inhibition of CXCR4 and indicates that they also need to be in the correct orientation and/or close proximity.

It is notable that CXCR4 can form complexes with membrane proteoglycans^44,45^ and taken together this leads us to propose two potential modes of inhibition by CXCL17: 1) A ‘bridge formation’ where CXCL17 simultaneously binds to CXCR4 and a GAG in close proximity to the receptor (Figure 5f) or 2) a result of clustering of GAGs induced by CXCL17 in the vicinity of CXCR4 causing steric hindrance and reduced access of CXCL12 to the receptor (Figure 5g). In the first mechanism, elements of the GAG sidechain must contribute to the binding of CXCL17 to CXCR4. While we recognise that for most chemokines simultaneous binding to GAGs and chemokine receptors is unlikely due to the overlap of chemokine binding sites and/or steric hinderance^32^, potential exceptions do exist; for example, for CCL21 and CXCL12γ that have extended C-terminal tails enriched in basic residues^21^. As the putative GAG binding domains of CXCL17 are positioned toward its C-terminus, similar to the situation with CCL21 and CXCL12γ, dual receptor-GAG binding may be structurally permitted. In support of option (2), chemokines are known to induce clustering of GAGs/proteoglycans^21^. Therefore, CXCL17-mediated clustering and stiffening of membrane proteoglycan complexes around CXCR4 could hinder the access of CXCL12 to the extracellular domains of CXCR4. This may also explain why CXCL17 did not interfere with the binding of a small molecule fluorescent derivative of IT1t, which binds within a transmembrane pocket of CXCR4^28^ and may more readily penetrate through clustered proteoglycans.

While the exact mechanism of CXCL17-mediated inhibition requires further investigation, it is worth noting that the extracellular domains of CXCR4 are highly acidic and inhibition by highly basic peptides is a known phenomenon^46^. Positively charged CXCR4 inhibitors include the synthetic peptides TT22 (18-amino acids of which 8 are positively charged), ALX40-4C (nonapeptide of 9 arginines), T134 and T140 (argine/lysine rich peptides)^47^ as well as the viral TAT protein^46^ and the endogenous cationic antimicrobial peptide cathelicidin LL37^23^. While here CXCL17 and the GAG inhibitors protamine sulfate and surfen are all positively charged, the fact that inhibition of CXCR4 only occurs in intact cells indicates a unique mechanism of action that differentiates these molecules from those previously described. Further investigation of this mechanism may yield a novel approach to therapeutically target CXCR4.

CXCL17 has been reported to facilitate angiogenesis principally via secretion of pro-angiogenic factors such as VEGF-A and b-FGF^6,37^. Here binding of CXCL17 to NRP1 represents a direct interaction with the prototypical VEGF-A angiogenic pathway. NRP1 acts as a co-receptor that selectively potentiates VEGFR2 signaling, a key driver of angiogenesis, and has been reported to directly promote angiogenesis^34^. While we did not directly assess the effect on VEGF-A mediated signaling or angiogenesis, CXCL17-mediated angiogenesis via interactions with NRP1 is plausible and should be examined further. Furthermore, the potential for interplay between chemokines and receptor tyrosine kinases (RTKs) due to potential overlapping glycosaminoglycan binding is an intriguing finding, particularly since some RTKs require formation of an obligate RTK-GAG complex for ligand binding, for example FGF-2 binding to FGFR^48^.

In summary, we demonstrate that CXCL17 inhibits CXCR4 and does so potentially via a glycosaminoglycan-containing accessory protein. These data suggest a unique mechanism of action that could be targeted to therapeutically inhibit CXCR4 in a cell type specific manner. Additionally, we confirm that GPR35 does not play a role in the function of CXCL17 and furthermore demonstrate CXCL17 has the potential to modulate VEGF signalling pathways via direct interactions with the VEGFR2 co-receptor NRP1. Together, the identification of CXCR4 and NRP1 as new molecular targets is a substantial contribution to our understanding of CXCL17 and may provide important insights into the patho/physiological roles of CXCL17.

## Methods

### Materials

AMD3100 was purchased from Selleckchem (USA) or Sigma-Aldrich (Australia). Unlabeled chemokine ligands were purchased from Preprotech (USA) except for 24Leu-CXCL17 which was purchased from R&D Systems and 22Ser-CXCL17 which was purchased from Sapphire Bioscience (Australia). CXCL12-AF647 was purchased from Almac (United Kingdom). N,N’-Dicyclohexylcarbamimidothioic acid (5,6-dihydro-6,6-dimethylimidazo[2,1-b]thiazol-3-yl)methyl ester dihydrochloride (IT1t) was purchased from Tocris (United Kingdom). Bovine serum albumin, Isoprenaline, Forskolin, Heparan sulfate sodium salt, saponin, surfen hydrate, protamine sulfate and zaprinast were purchased from Sigma-Aldrich (Australia/United Kingdom). Furimazine, caged-furimazine (Endurazine), EnduRen, GloSensor cAMP reagent and purified LgBiT NLuc fragments were from Promega Corporation (USA). SNAPTag AlexaFluor 488 membrane-impermeant substrate was from New England Biolabs. VEGF_165a_ was purchased from R&D Systems and synthesis of fluorescent VEGF_165a_-tetramethylrhodamine (TMR) has been described previously^36^.

IT1t-BY630-650 ((6,6-Dimethyl-5,6-dihydroimidazo[2,1-b]thiazol-3-yl)methyl (E)-N-cyclohexyl-N’-((1r,4r)-4-(6-(2-(4-((E)-2-(5,5-difluoro-7-(thiophen-2-yl)-5H-4λ4,5λ4-dipyrrolo[1,2-c:2’,1’-f][1,3,2]diazaborinin-3-yl)vinyl)phenoxy)acetamido)hexanamido)cyclo-hexyl)carbamimidothioate) was synthesised in the University of Nottingham. Purity of the compound was confirmed to be >95% by analytical HPLC. HRMS (ESI-TOF) calculated for C_50_H_61_B_1_F_2_N_8_O_3_S_3_ [M + H]^+^: 967.4168 found: 967.4238. Full synthetic details will be communicated shortly (manuscript in preparation).

AMD3100 was dissolved in water, unlabeled chemokines and CXCL12-AF647 were dissolved as per the manufacturer’s instructions. VEGF_165_a was dissolved in phosphate buffered saline (PBS) containing 0.1% bovine serum albumin (BSA; Sigma Aldrich). IT1t and Zaprinast were dissolved in DMSO to a concentration of 10 mM. All further dilutions were performed in assay buffer containing 0.1% BSA.

### Molecular biology and construct sources

Expression cDNA constructs encoding untagged chemokine receptors, β2-adrenoceptor or GPR35 were purchased from the cDNA Resource Center. To generate the pcDNA3.1(+) mammalian expression vector encoding for GPR35 tagged on the C-terminus with NLuc, the stop codon was first removed by site directed mutagenesis. The primers used were forward 5’-CCTCTAGACTCGAGTGCGGCGAGGGTCACGCA-3’ and reverse 5’-TGCGTGACCCTCGCCGCACTCGAGTCTAGAGG-3’. Nluc was then sub-cloned into the vector from existing constructs using the restriction enzymes XhoI and ApaI. Generation of pcDNA3.1 (+) neo expression constructs encoding NLuc/CXCR4, HiBiT/CXCR4^49^, SNAP/CXCR4^29^, NLuc/VEGFR2^36^ and NLuc/NRP^35^ have been described previously, as have pcDNA3 expression constructs encoding CXCR4/Rluc8^50^, CXCR4/N luc^51^, β-arrestin2/Venus^52^. Venus/G_γ2_ was a kind gift from Dr. Martina Kocan. Gα_i1_/NLuc was generated from the Gαi1/Rluc8 construct reported previously^53^. The N119S CXCR4 mutant was generated by site-directed mutagenesis. The primers used were forward 5’-CATCTACACAGTCAGCCTCTACAGCAGTG-3’ and reverse 5’-CACTGCTGTAGAGGCTGACTGTGTAGATG-3’.

### Cell culture

HEK293F/T (Life Technologies), HEK293G (Promega), HeLa (gift from Steve Briddon, University of Nottingham) and Cos-7 (ATCC) cells were maintained in Dulbecco’s Modified Eagle’s Medium (Sigma Aldrich or Thermo Fisher Scientific) supplemented with 10 % fetal calf serum at 37°C/5% CO2. Stable transfections were performed using FuGENE (Promega, USA) according to the manufacturer’s instructions. Generation of HEK293 cell lines stably-expressing Nluc/CXCR4, HiBiT/CXCR4 or SNAP/CXCR4, or generation of HEK293 or HeLa cells that were genome-edited to express NLuc/CXCR4 or CXCL12-HiBiT under endogenous promotion, have been described previously^29,49^. To generate cells for use in the GloSensor cAMP assay, HEK293G cells stably-expressing the GloSensor cAMP biosensor were transfected with a pcDNA3.1 (+) neo expression vector encoding SNAP/CXCR4 and subsequently selected for incorporation of the transgene using G418 (ThermoFisher).

### BRET β-arrestin2 recruitment assays

HEK293F/T cells were maintained in DMEM containing 0.3 mg/ml glutamine, 100 IU/ml penicillin and 100 μg/ml streptomycin supplemented with 10% FBS (Bovogen) and 400-μg/mL geneticin. Cells were harvested when cells reached 70-80% confluency using phosphate-buffered saline (PBS) and 0.05% trypsin-EDTA and seeded in a 6 well plate at 300,000 cells/well. 24 hours later cells were transiently transfected with pcDNA3 mammalian expression vectors encoding CXCR4/Rluc8 (100 ng per well) and β-arrestin2/Venus (300 ng per well) using FuGENE and incubated at 37°C/5% CO_2_ for 24 h. Cells were harvested into white 96-well plates at 100,000 cells/well in phenol-red free DMEM containing 25 mM HEPES, 0.3 mg/ml glutamine, 100 IU/ml penicillin and 100 μg/ml streptomycin supplemented with 5% FCS (phenol-red free DMEM) and incubated at 37°C/5% CO_2_ for 24 h. On the day of assay, culture medium was removed and cells incubated with EnduRen (30 μM) in pre-warmed Hanks-buffered saline solution without calcium or magnesium (Hanks buffer, Thermo Fisher Scientific) for 2 hours at 37 °C. Real-time BRET measurements were taken at 37 °C using a LUMIstar. In all experiments, initial measurements were taken to establish the baseline BRET ratio before CXCL12 (10 nM) or CXCL17 1 pM – 1 μM) was added to the cells. In a subset of experiments, vehicle or CXCL12 (10 nM) was added and the BRET allowed to plateau before CXCL17 (100 nM) was added to the cells. Filtered light emissions were sequentially measured at 475 nm (30 nm bandpass) and 535 nm (30 nm bandpass). The corrected BRET signal was calculated by subtracting the ratio of the acceptor 535/30 nm emission over the donor 475/30 nm emission for a vehicle-treated cell sample from the same ratio for a sample treated with agonist. In this calculation, the vehicle-treated cell sample represents the background, eliminating the requirement for measuring a donor-only control sample. Points represent the maximal change in the BRET ratio following ligand addition from the kinetic read. In a subset of experiments, HEK293 cells where transfected with cDNA encoding CXCR1/RLuc8, CCR5/RLuc8 or β_2_-adrenoceptor/RLuc8 (100 ng per well) and β-arrestin2/Venus (300 ng per well), and following establishment of the baseline BRET ratio, cells were incubated with either CXCL17 (300 nM) or CXCL8 (10 nM), CCL5 (10 nM) or Isoprenaline (100 μM) respectively.

To investigate β-arrestin/Venus recruitment to GPR35/NLuc, 300,000 cells in a 6 well plate were transfected with pcDNA3 mammalian expression vectors encoding GPR35/NLuc (25 ng per well) and β-arrestin2/Venus (300 ng per well), then seeded at 30,000 cells/well in a 96 well plate 24 hours later. On the day of assay, cells were washed and incubated with Hanks buffer for 1 hour at 37 °C before furimazine (10 μM) was added. Cells were incubated for a further 5 mins at 37 °C before real-time BRET measurements were taken at 37 °C using a LUMIstar as described above. Following establishment of the baseline BRET ratio, cells were treated with vehicle or CXCL17 (100 nM) or Zaprinast (100 μM).

To establish the pIC_50_ of CXCL17 in figure 2c, HEK293 cells were maintained in Dulbecco’s Modified Eagle’s Medium (Sigma Aldrich) supplemented with 10 % fetal calf serum at 37°C/5% CO_2_. Cells were passaged or harvested when cells reached 70-80% confluency using Phosphate-Buffered Saline (PBS, Sigma Aldrich) and trypsin (0.25% w/v in versene; Sigma Aldrich). Cells were seeded at 300,000 cells/well in a 6 well plate and incubated for 24h at 37°C/5%CO_2_ before being transfected with CXCR4/NLuc (25 ng per well) and β-arrestin2/Venus (300 ng per well) using FuGENE. Cells were seeded into poly-D-lysine coated white flat bottom 96 well plates at 40,000 cells/well and incubated for 24h at 37°C/5% CO_2_. On the day of the assay, cells were washed and incubated with pre-warmed HEPES-Buffered Salt Solution (HBSS; 25mM HEPES, 10mM glucose, 146mM NaCl, 5mM KCl, 1mM MgSO4, 2mM sodium pyruvate, 1.3mM CaCl2, 1.8g/L glucose; pH 7.2) supplemented with 0.1% BSA containing CXCL17 (0.3 nM – 1 μM) or AMD3100 (10 pM – 300 nM) for 1 hour at 37°C. Following incubation with furimazine (10 μM) for 5 minutes before sequential filtered light emissions were taken using a PHERAStar FS plate reader using 475 nm (30 nm bandpass) and 535 nm (30 nm bandpass) filters. Following establishment of the baseline BRET ratio, vehicle or CXCL12 (10 nM) was added to the cells before further filtered light emissions were taken. The corrected BRET signal was calculated by subtracting the ratio of the acceptor 535/30 nm emission over the donor 475/30 nm emission for a vehicle-treated cell sample from the same ratio for a sample treated with agonist.

### G protein activation assays

HEK293 or Cos-7 cells were maintained in DMEM containing 0.3 mg/ml glutamine, 100 IU/ml penicillin and 100 μg/ml streptomycin supplemented with 10% FBS (Bovogen) and 400-μg/mL geneticin. Cells were seeded in a 6 well plate at 300,000 cells/well and 24 hours later cells were transiently transfected with pcDNA3 mammalian expression vectors encoding Gα_i1_/Nluc (50 ng/well) and Venus/G_γ2_ (100 ng/well) as well as empty pcDNA3 or GPR35 (50 ng/well) with or without cDNA encoding a chemokine receptor (CCR1-CCR10 or CXCR1-CXCR6, 125 ng/well) or alternatively cells were transfected with Gα_i1_/Nluc (50 ng/well), Venus/G_γ2_ (100 ng/well) as well as chemokine receptor (CCR1-CCR10 or CXCR1-CXCR6 or N119S CXCR4 mutant, 100 ng/well) using FuGENE and incubated at 37°C/5% CO_2_ for 24 h. Cells were then harvested and were seeded into poly-L-lysine coated white flat bottom 96 well plates in phenol-red free DMEM at 50,000 cells/well and incubated for 24h at 37°C/5% CO_2_. On the day of the assay, medium was aspirated and the cells were incubated with pre-warmed Hanks buffer supplemented with 0.1% BSA containing Endurazine (10 μM) for 2 hours at 37°C. Sequential filtered light emissions were then recorded using a LUMIstar plate reader using 475 nm (30 nm bandpass) and 535 nm (30 nm bandpass) filters to establish a baseline BRET ratio. Cells were then treated with ligands and the BRET ratio continuously monitored. In assays investigating GPR35 only, cells were treated with Zaprinast (100 μM, or 10 nM – 300 μM for concentration response curves) or CXCL17 (100 nM). Where GPR35 and a chemokine receptor were co-expressed, cells were stimulated with CXCL17 (30 nM) or 30 nM of CCL3, CCL2, CCL13, CCL22, CCL4, CCL20, CCL19, CCL1, CCL25, CCL27, CXCL8, CXCL8, CXCL11, CXCL12, CXCL13 and CXCL16 for CCR1-CCR10 and CXCR1-6 respectively. HEK293 cells expressing CXCR4 were treated with CXCL12 (1 nM), CXCL17 (300 nM), or CXCL12 (1 nM) and CXCL17 (300 nM). Cells expressing CXCR4 N119S or wildtype CXCR4 in the parallel control were treated with AMD3100 (10 μM), CXCL12 (10 nM) or CXCL17 (100 nM). Cos-7 cells expressing CXCR4 were treated with CXCL12 (10 nM), CXCL17 (300 nM) or CXCL12 (10 nM) and CXCL17 (300 nM). In HEK293 cells where CCR1-CCR10 and CXCR1-3, 5 and 6 were expressed, cells were treated with CXCL17 (300 nM) or a submaximal concentration of chemokine CCL3 (0.3 nM), CCL2 (0.3 nM), CCL13 (1 nM), CCL22 (0.3 nM), CCL4 (30 nM), CCL20 (3 nM), CCL19 (1 nM), CCL1 (10 nM), CCL25 (10 nM), CCL27 (10 nM), CXCL8 (0.3 nM), CXCL8 (1 nM), CXCL11 (1 nM), CXCL13 (3 nM) and CXCL16 (1 nM) for CCR1-10 and CXCR1-3, 5 and 6 respectively in the absence or presence of CXCL17 (300 nM). In these experiments’ cells were preincubated with CXCL17 for 15 mins prior to establishing the baseline BRET ratio. Corrected BRET ratios were calculated by subtracting the ratio of the 535 nm emission (acceptor) by the 475 nm emission (donor) for a vehicle-treated cell sample from the same ratio for a sample treated with agonist. For concentration-response curves or bar graphs, data represent the maximal change in the corrected BRET ratio.

To establish the pIC_50_ of CXCL17 in figure 2c, HEK293 cells were maintained in Dulbecco’s Modified Eagle’s Medium (Sigma Aldrich) supplemented with 10% fetal calf serum at 37°C/5% CO_2_. Cells were harvested when they reached 70-80% confluency using Phosphate Buffered Saline (PBS, Sigma Aldrich) and trypsin (0.25% w/v in versene; Sigma Aldrich) and seeded at 300,000 cells/well in a 6 well plate and incubated for 24h at 37°C/5% CO2 before being transfected with CXCR4 (10 ng per well), Gα_i1_/Nluc (50 ng/well) and Venus/G_γ2_ (100 ng/well) using FuGENE. 24 hours later, cells were harvested and seeded into poly-D-lysine coated white flat bottom 96 well plates at 40,000 cells/well and incubated for 24h at 37°C/5% CO_2_. On the day of the assay, cells were washed and incubated with pre-warmed HBSS supplemented with 0.1% BSA containing CXCL17 (0.3 nM – 1 μM) or AMD3100 (10 pM – 300 nM) for 30 minutes at 37°C. Cells were then incubated with furimazine (10 μM) for 10 minutes before filtered light emissions were taken using a PHERAStar FS plate reader using 475 nm (30 nm bandpass) and 535 nm (30 nm bandpass) filters. Following establishment of the baseline BRET ratio, vehicle or CXCL12 (0.1 nM) was added to the cells before further filtered light emissions were taken. The corrected BRET signal was calculated by subtracting the ratio of the acceptor 535/30 nm emission over the donor 475/30 nm emission for a vehicle-treated cell sample from the same ratio for a sample treated with agonist.

### GloSensor cAMP assay

GloSensor cAMP assay was performed according to the manufacturer’s instructions. Briefly, HEK293G cells stably expressing SNAP/CXCR4 were seeded into poly-D-lysine coated white flat bottom 96 well plates at 30,000 cells/well and incubated for 24h at 37°C/5% CO_2_. On the day of the assay, cells were washed then incubated with pre-warmed HBSS containing 6% GloSensor cAMP reagent and 0.1% BSA for 1.5 hours at 37°C. Cells were then pre-incubated with or without AMD3100 (1μM) or CXCL17 (1 μM) for 30 minutes before total light emissions were measured on a PHERAStar FS plate reader to establish baseline luminescence. Forskolin (30 μM) in the absence or presence of CXCL12 (1 nM) was then added and total luminescence measured for a further 1 hour. Data represents % of the forskolin (30 μM) mediated cAMP production and was calculated from the area under the curve following ligand addition.

### NanoBRET ligand binding assays

HEK293T cells stably expressing NLuc/CXCR4, HiBiT/CXCR4 or HEK293F/T as well as HeLa cells expressing genome-edited NLuc/CXCR4 were maintained in Dulbecco’s Modified Eagle’s Medium (Sigma Aldrich) supplemented with 10% fetal calf serum at 37°C/5% CO_2_. Cells were passaged or harvested when they reached 70-80% confluency using Phosphate Buffered Saline (PBS, Sigma Aldrich) and trypsin (0.25% w/v in versene; Sigma Aldrich). Membrane preparations expressing genome-edited or stably-expressed NLuc/CXCR4 were made as described previously^54^.

For competition binding experiments, cells expressing NLuc/CXCR4 were seeded into poly-D-lysine coated white flat bottom 96 well plates at 30,000 cells/well and incubated for 24h at 37°C/5% CO_2_. On the day of the assay, cells were washed and incubated with pre-warmed HBSS supplemented with 0.1% BSA. For assays using membrane preparations expressing NLuc/CXCR4 from genome-edited or stably transfected HEK293 cells, 10 μg membrane protein diluted in HBSS supplemented with 0.1% BSA was loaded into each well. Cells or membranes were then incubated with CXCL12-AF647 (12.5 nM) or IT1t-BY630/650 (100 nM) in the absence or presence of AMD3100 (1 pM – 10 μM), CXCL4 (1 μM), CXCL12 (0.1 pM – 1 μM), CXCL17 (1pM – 1 μM), IT1t (1 pM – 10 μM), Surfen (10 pM – 10 μM) or protamine sulfate (1nM – 10 μM) for 1 hour at 37°C. All experiments testing an unknown inhibitor of CXCR4 (CXCL17, Surfen or Protamine sulfate) where performed with a known CXCR4 ligand (AMD3100, CXCL12, IT1t) as a control. Following incubation, furimazine (10 μM) was added and plates equilibrated for 5 minutes at room temperature before sequential filtered light emissions were taken using a PHERAStar FS plate reader using 460nm (80nm bandpass) and >610nm (longpass) filters. BRET ratios were calculated by dividing the 610nm emission (acceptor) by the 460nm emission (donor). For cells expressing HiBiT/CXCR4, experiments were performed as described above but following ligand incubation both furimazine (10 μM) and purified LgBiT (10 nM) were added to generate luminescence.

In subsets of experiments the NanoBRET competition binding assays were performed as described above but with the following modifications: In figure 3d, to determine the effect of membrane permeabilization with saponin, 10 μg membrane protein diluted in HBSS supplemented with 0.1% BSA was loaded into each well containing 0.25 mg/mL saponin and membranes incubated with CXCL12-AF647 (12.5 nM) in the absence or presence of AMD3100 (1 μM) or CXCL17 (300 nM) for 1 hour at 37°C. In figure 3e, to determine if cellular proteases were required to convert CXCL17 into a more active species, wildtype HEK293 cells were seeded into poly-D-lysine coated white flat bottom 96 well plates at 30,000 cells/well and incubated for 24h at 37°C/5% CO_2_. On the day of the assay, cells were washed and 10 μg membrane protein diluted in HBSS supplemented with 0.1% BSA loaded into each well containing cells and incubated with CXCL12-AF647 (12.5 nM) in the absence or presence of increasing concentrations of AMD3100 (1 pM – 10 μM) or CXCL17 (100 pM – 1 μM), for 1 hour at 37°C. In Supplementary figure 7b, to investigate the effect of exogenous heparan sulfate on CXCL17, on the day of the assay HEK293 cells expressing NLuc/CXCR4 were washed and incubated with prewarmed HBSS supplemented with 0.1% BSA before AMD3100 (1 μM) or CXCL17 (300 nM) that had been preincubated with or without heparan sulfate (30 μg/ml) for 1 hour was added to the cells. CXCL12-AF647 (12.5 nM) was then added to each well and incubated for 30 mins at 37°C before the plate was read.

For competition binding experiments expressing NLuc/VEGFR2 or NLuc/NRP1, wildtype HEK293 cells were seeded into poly-D-lysine coated white flat bottom 96 well plates at 10,000 cells/well and incubated for 24h at 37°C/5% CO_2_. Cells were then transiently-transfected with 100 ng/well cDNA encoding NLuc/NRP1 or NLuc/VEGFR2 using FuGENE (Promega, USA) according to the manufacturer’s instructions and incubated for a further 24h at 37°C/5% CO_2_. On the day of the assay, cells were washed and incubated with pre-warmed HBSS supplemented with 0.1% BSA. Cells were then incubated with VEGF_165_a-TMR (3 nM) in the absence or presence of CXCL12 (1 μM), CXCL17 (300 nM), Surfen (30 μM) or VEGF165a (30 μM) for 1 hour at 37°C. Following incubation, furimazine (10 μM) was added and plates equilibrated for 5 minutes at room temperature before sequential filtered light emissions were measured using a PHERAStar FS plate reader using 460nm (80nm bandpass) and >610nm (longpass) filters. BRET ratios were calculated by dividing the 610nm emission (acceptor) by the 460nm emission (donor).

### Determination of IT1t-BY630-650 affinity by saturation NanoBRET saturation ligand binding

Genome-edited cells expressing NLuc/CXCR4 were seeded into poly-D-lysine coated white flat bottom 96 well plates at 30,000 cells/well and incubated for 24h at 37°C/5% CO_2_. On the day of the assay, cells were washed and incubated with pre-warmed HBSS supplemented with 0.1% BSA and incubated with increasing concentrations of IT1t-BY630/650 in the absence or presence of IT1t (10 μM) for 1 hour at 37°C. Furimazine (10 μM) was added, and plates equilibrated for 5 minutes at room temperature before sequential filtered light emissions were taken using a PHERAStar FS plate reader using 460nm (80nm bandpass) and >610nm (longpass) filters. BRET ratios were calculated by dividing the 610nm emission (acceptor) by the 460nm emission (donor).

### Nano luciferase complementation internalisation assays

HEK293 cells stably expressing HiBiT/CXCR4 were seeded into poly-D-lysine coated white flat bottom 96 well plates at 30,000 cells/well and incubated for 24h at 37°C/5% CO_2_. On the day of the assay, cells were washed and incubated with pre-warmed HBSS supplemented with 0.1% BSA. A concentration-response curve was generated by incubating cells with CXCL17 (10 pM – 1 uM) for 60 minutes at 37°C. Following ligand incubation, furimazine (10 μM) and purified LgBiT (10 nM) were added and plates incubated for 5 minutes before total light emissions were continuously measured using a PHERAStar FS plate reader with the concentration-response curves representing the luminescence after 30 minutes.

### Internalisation by fluorescent microscopy

HEK293G cells stably expressing SNAP/CXCR4 were seeded into poly-D-lysine coated 8-well plates at 10,000 cells/well and incubated for 24h at 37°C/5% CO_2_. On the day of the assay, cells were incubated with 0.5 μM membrane impermeant SNAP-tag AF488 for 30 minutes at 37°C/5% CO_2_ prepared in serum-free DMEM. Cells were then washed three times with HBSS/0.1% BSA and incubated with CXCL12 (100 nM), CXCL17 (300 nM) or vehicle (HBSS/0.1% BSA) for 1 hour in the dark at 37°C. Cells were imaged live at 37°C using a Zeiss LSM710 fitted with a 63x Plan Apochromat oil objective (1.4NA) using a Argon488 (AlexaFluor488; 496-574nm band pass; 2% power) with a 488/561/633 beamsplitter using a pinhole diameter of 1 Airy unit. All images were taken at 1024×1024 pixels per frame with 8 averages (zoom 1).

### CXCL12-HiBiT NanoBRET competition ligand binding assays

HEK293 cells expressing genome-edited CXCL12-HiBiT were seeded in 6 well plates at 300,000 cells per well and incubated for 24h at 37°C/5% CO_2_. Cells were then transfected with 500 ng/well pcDNA3.1 (neo) plasmid encoding SNAP/CXCR4 and incubated for a further 24h before seeding into poly-D-lysine coated white flat bottom 96 well plates at 60,000 cells/well and incubated for 24h at 37°C/5% CO_2_. On the day of the assay, cells were incubated with 0.25 μM membrane impermeant SNAP-tag AF488 for 1h at 37°C/5% CO_2_ prepared in serum-free DMEM. After incubation, cells were washed 3 times with pre-warmed HBSS supplemented with 0.1% BSA and incubated with purified 30 nM purified LgBiT in the absence or presence of AMD3100 (1 pM – 10 μM), Surfen (3 nM – 30 μM) or CXCL17 (100 pM – 1 μM) for 2h at 37°C. Following ligand incubation, 10 μM furimazine was added and plates equilibrated for 5 mins at room temperature. To determine basal BRET, SNAP-tagAF488 was omitted during labelling. Sequential filtered light emissions were recorded using a PHERAStar FS plate reader using 475 nm (30 nm bandpass) and 535 nm (30 nm bandpass) filters. BRET ratios were calculated by dividing the 535 nm emission (acceptor) by the 475 nm emission (donor).

### CXCL17 conservation analysis

Conservation analysis was performed by WebLogo^55^ using the multiple sequence alignment of forty-five 119 amino acid CXCL17 orthologues obtained from UniProt^56^ (Supplementary Table 1).

### Data presentation and statistical analysis

Due to differences in plate reader sensitivity and expression differences between cell lines/assays, raw BRET ratios cannot be compared as a measure of BRET efficacy between figures as optimised plate reader emission gains were used to ensure sufficient sensitivity and/or measurements acquired did not saturate the detector. In general, BRET ratios were calculated by dividing the acceptor emission by the donor emission. Calculation of baseline-corrected BRET ratios or luminescence values are described in the methods for each assay configuration. Where results reported are taken from kinetic reads, points or bars are the maximum ligand-induced change in the BRET ratio.

Prism 8 software was used to analyse ligand-binding curves. For NanoBRET receptor-ligand saturation binding assays, total and non-specific saturation binding curves were simultaneously fitted using the following equation:

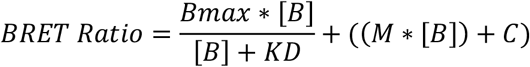

where Bmax is the maximal response, [B] is the concentration of fluorescent ligand in nM, KD is the equilibrium dissociation constant in nM, M is the slope of the non-specific binding component and C is the intercept with the Y-axis.

Concentration-response data were fitted using the following equation:

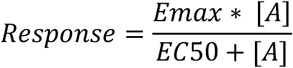

Where Emax is the maximum response, EC_50_ is the concentration of agonist required to produce 50% of the maximal response and [A] is the agonist concentration. pEC_50_ calculated from maximal change in BRET.

Competition binding or response data were fitted using the following equation:

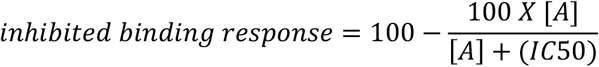

where [A] is the concentration of competing ligand, IC_50_ is the molar concentration of this competing ligand required to inhibit 50% of the specific response or binding.

In binding studies, the Cheng-Prusoff equation was used to correct fitted IC_50_ values to K_i_ values:

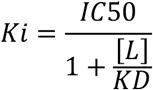

Where [L] is the concentration of fluorescent ligand in nM and K_D_ is the dissociation constant of fluorescent ligand in nM. K_D_ values calculated from saturation binding experiments^49^.

Statistical analysis was performed using Prism 7 or 8 software (GraphPad, San Diego, USA) using one or two-way ANOVA with an appropriate multiple comparison’s test where required or paired t-test. Specific statistical tests used are indicated in the figure legends and were performed on the mean data of individual experiments (n) also indicated in the figure legends. A p-value <0.05 was considered statistically significant.

## Acknowledgements

This work was supported by a MRC grant number MR/N020081/1, a European Union’s Horizon 2020 MSCA Programme grant (ONCORNET, agreement 641833). C.W.W. is supported by an NHMRC CJ Martin Fellowship (1088334) and by a UWA fellowship support grant. L.E.K. is supported by a University of Nottingham Anne McLaren Research Fellowship. N.D. is supported by an Australian Government Research Training Program Scholarship and a University of Western Australia Baillieu Research Scholarship. The authors would like to thank Heng See for technical assistance with preliminary experiments and Dr. Brigit Caspar for generation of CXCR4 expression plasmids and cell lines. Figures 5 f-g were created using BioRender.

## Author contributions

C.W.W conceived the study. C.W.W., L.E.K., M.J.S and S.D generated reagents. C.W.W., L.E.K., N.D. and R.A. conducted the experiments. C.W.W and L.E.K. performed the data analysis. C.W.W., L.E.K., K.D.G.P., and S.J.H. wrote or contributed to the writing of the manuscript.

