## Supplementary Information for "CXCL17 is an endogenous inhibitor of CXCR4 via a novel mechanism of action"

**Supplementary Figure 1**

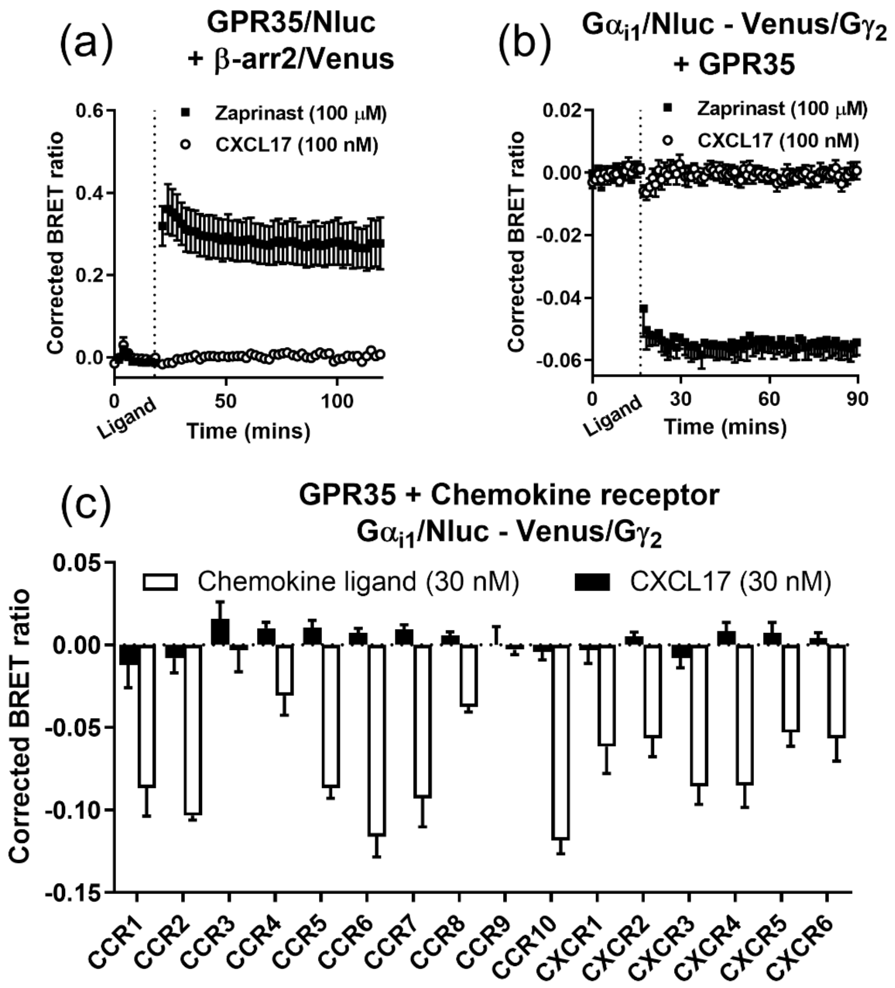


**Supplementary Figure 1: CXCL17 does not activate GPR35. (a)** HEK293 cells transiently transfected with GPR35/NLuc and β-arrestin2/Venus were stimulated with zaprinast (100 μM, black squares) or CXCL17 (100 nM, white circles). **(b)** HEK293 cells transiently transfected with GPR35, $G\alpha_{i1}$/NLuc and Venus/$G\gamma_2$ were stimulated with zaprinast (100 μM, black squares) or CXCL17 (100 nM, white circles). **(c)** Maximum change in BRET observed in HEK293 cells transiently-transfected with GPR35, $G\alpha_{i1}$/NLuc and Venus/$G\gamma_2$ plus CCR1-CCR10 or CXCR1-6 following application of the endogenous chemokine ligand (white bar, 30 nM: CCL3, CCL2, CCL13, CCL22, CCL4, CCL20, CCL19, CCL1, CCL25, CCL27, CXCL8, CXCL8 CXCL11, CXCL12, CXCL13 and CXCL16 respectively) or CXCL17 (black bar, 30 nM). For **a** and **b**, ligand was added following establishment of basal BRET and is indicated on the x-axis as *ligand.* Corrected BRET was calculated as described in *Methods.* Points or bars represent mean ± s.e.m. of three (**a** and **b**) or mean ± range of two (**c**) individual experiments performed in duplicate or triplicate.

**Supplementary Figure 2**

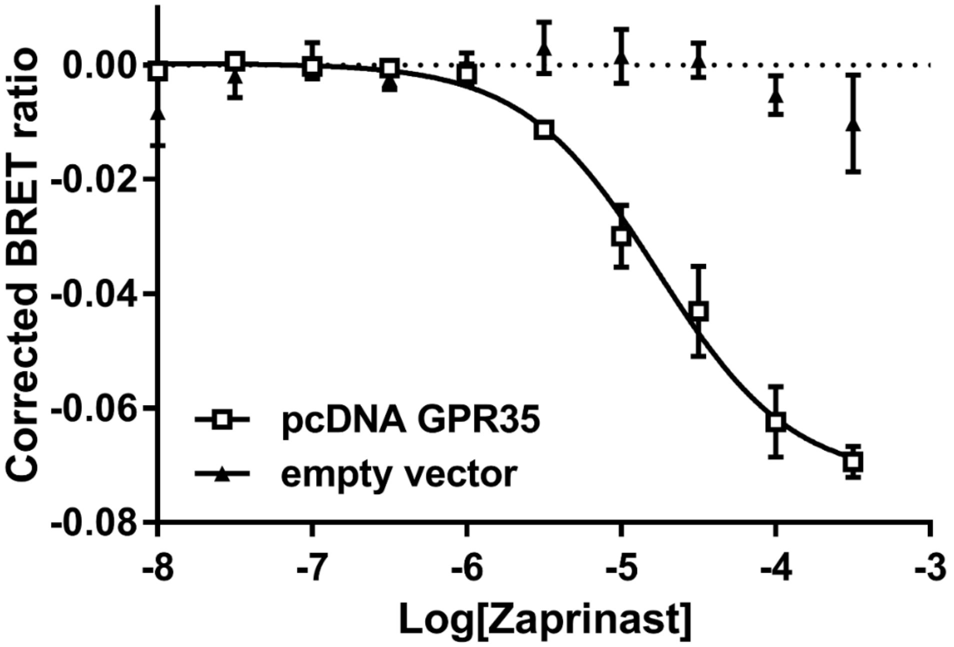


**Supplementary Figure 2: The $G_{\alpha i1}$/NLuc and Venus/$G_{\gamma 2}$ BRET assay measures GPR35 activation.** HEK293 cells transiently transfected with $G_{\alpha i1}$/NLuc and Venus/$G_{\gamma 2}$ along with plasmids encoding GPR35 (white squares, $pEC_{50} = 4.76 \pm 0.82$) or empty vector (black triangles) were stimulated with increasing concentrations of zaprinast (10 nM – 300 µM). Corrected BRET was calculated as described in Methods. Points represent mean ± s.e.m. of three individual experiments performed in triplicate.

**Supplementary Figure 3**

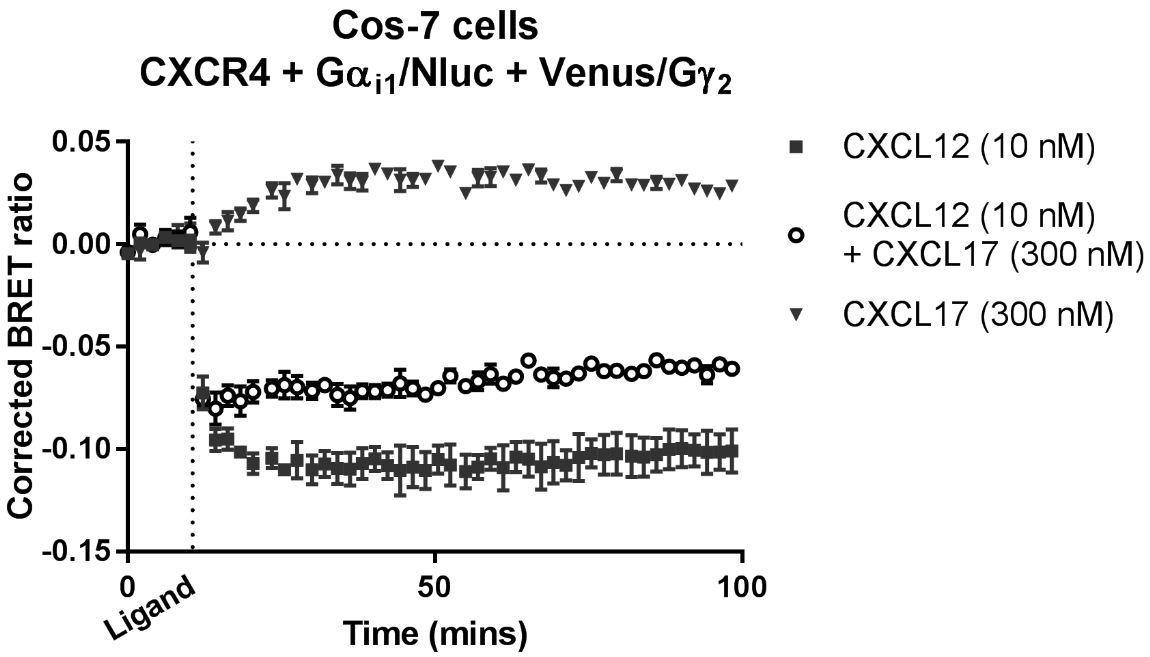


**Supplementary Figure 3: CXCL17 inhibits CXCR4-mediated G protein activation in Cos-7 cells**. Cos-7 cells transiently-transfected with Gαi1/NLuc, Venus/G$\gamma_2$ and CXCR4 were stimulated with CXCL12 (10 nM, black squares), CXCL12 (10 nM) and CXCL17 (300 nM, white circles) or CXCL17 (300 nM, black triangles). Corrected BRET was calculated as described in Methods. Points represent mean ± s.e.m. of three individual experiments performed in triplicate.

**Supplementary Figure 4**

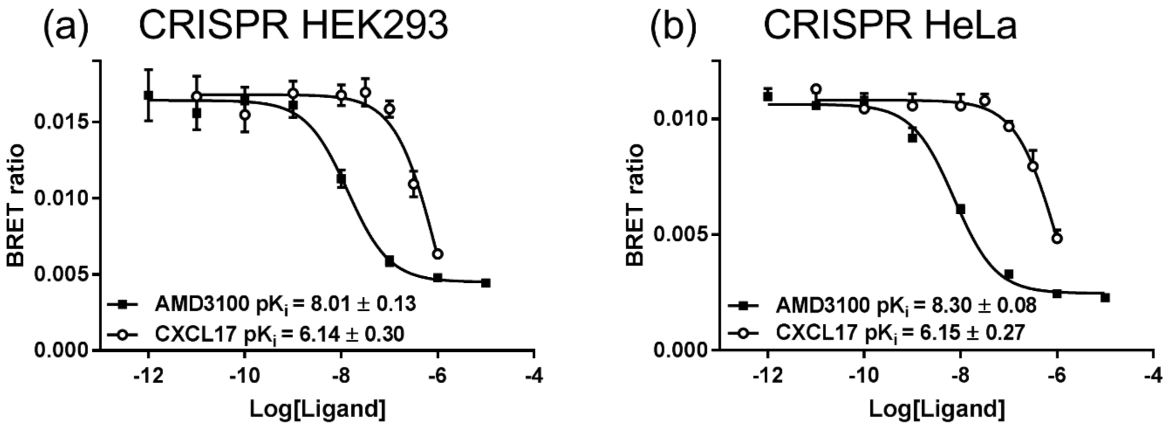


**Supplementary Figure 4: NanoBRET competition ligand binding at NLuc/CXCR4 expressed under endogenous promotion**. Displacement of 12.5 nM CXCL12-AF647 binding by AMD3100 (10 pM – 10 µM, black squares) or CXCL17 (100 pM – 1 µM, white circles) in live **(a)** HEK293 cells or **(b)** HeLa cells genome-edited to express NLuc/CXCR4. Points represent mean ± s.e.m. of four individual experiments performed in duplicate.

**Supplementary Figure 5**

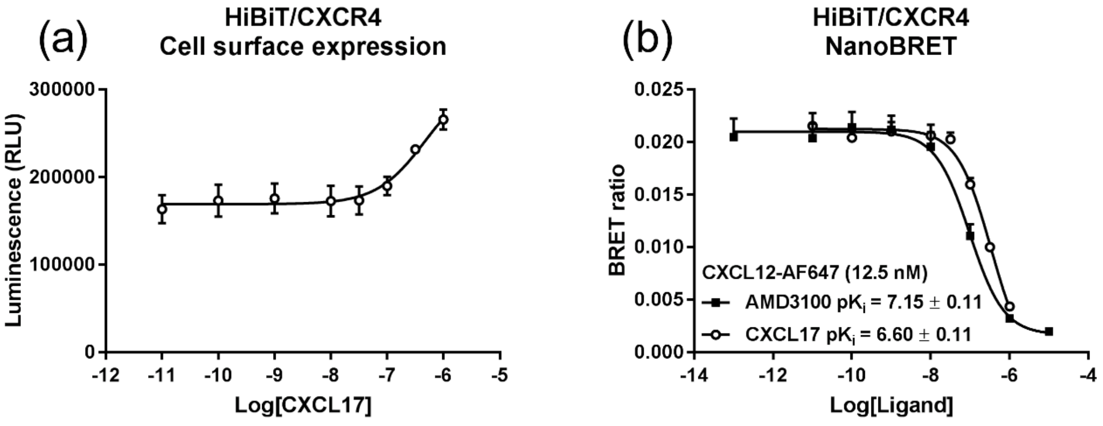


**Supplementary Figure 5: Effect of CXCL17 on CXCR4 cell surface expression and CXCL12-AF647 binding to plasma membrane-localised CXCR4.** **(a)** HEK293 cells stably-expressing HiBiT/CXCR4 were incubated with increasing concentrations of CXCL17 (10 pM - 1 µM) for 1h at 37°C and luminescence measured 30 min following addition of purified LgBiT (10 nM) and furimazine (10 µM). **(b)** NanoBRET competition ligand binding curves obtained in HEK293 cells stably-expressing HiBiT/CXCR4 incubated with CXCL12-AF647 (12.5 nM) and increasing concentrations of CXCL17 (10 pM - 1 µM, white circles) or AMD3100 (0.1 pM - 10 µM, black squares) for 1h at 37°C. To localise binding to the plasma membrane, NanoBRET was measured following the addition of purified LgBiT (10 nM) and furimazine (10 µM). Points represent mean ± s.e.m. of four **(a)** or five **(b)** individual experiments performed in duplicate.

**Supplementary Figure 6:**

**(a)**

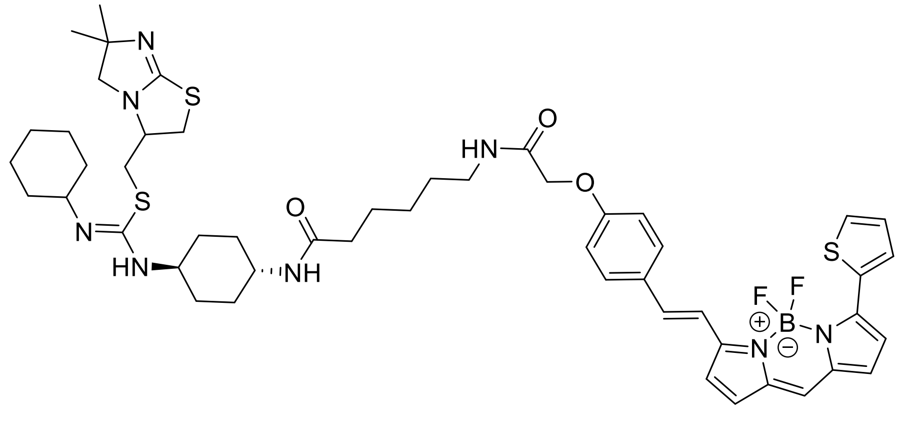


**(b)**

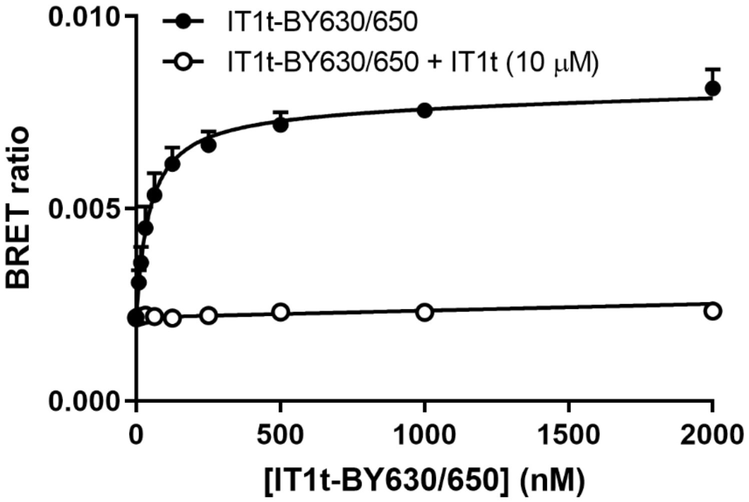


**Supplementary Figure 6: Determination of the binding affinity of IT1t-BY630/650 at NLuc/CXCR4 in HEK293 cells.** (a) Chemical structure of IT1t-BY630/650. (b) NanoBRET saturation ligand binding curves obtained in HEK293 cells expressing genome-edited NLuc/CXCR4. Cells were incubated with increasing concentrations of IT1t-BY630/650 in the absence (black circles) or presence (white circles) of IT1t (10 μM) for 1 h at 37°C. Data shown represent mean ± SEM of five independent experiments performed in duplicate

**Supplementary Figure 7:**

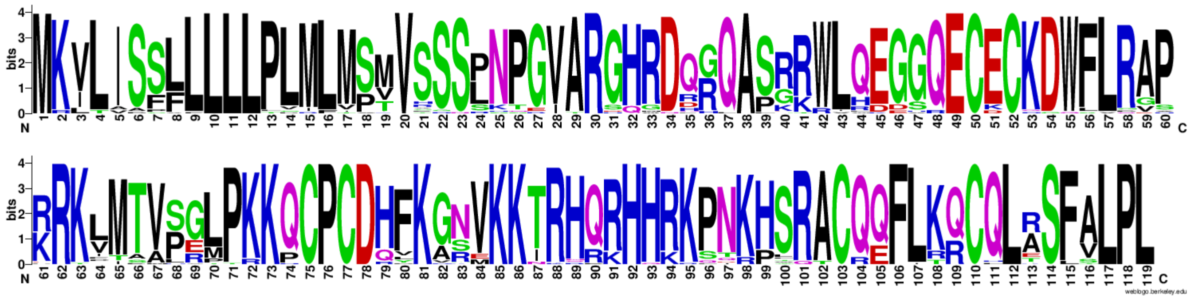

**Supplementary Figure 7: Conservation analysis of CXCL17.** 'Sequence logo' generated for CXCL17 based on the multiple sequence alignment of forty-five 119 amino acid CXCL17 orthologues (Supplementary Table 1). Logo was generated using WebLogo (Crooks et al., 2004). Putative GAG binding sites between amino acids 85-101 appear highly conserved. See Figure 5a for putative GAG domains.

**Supplementary Figure 8**

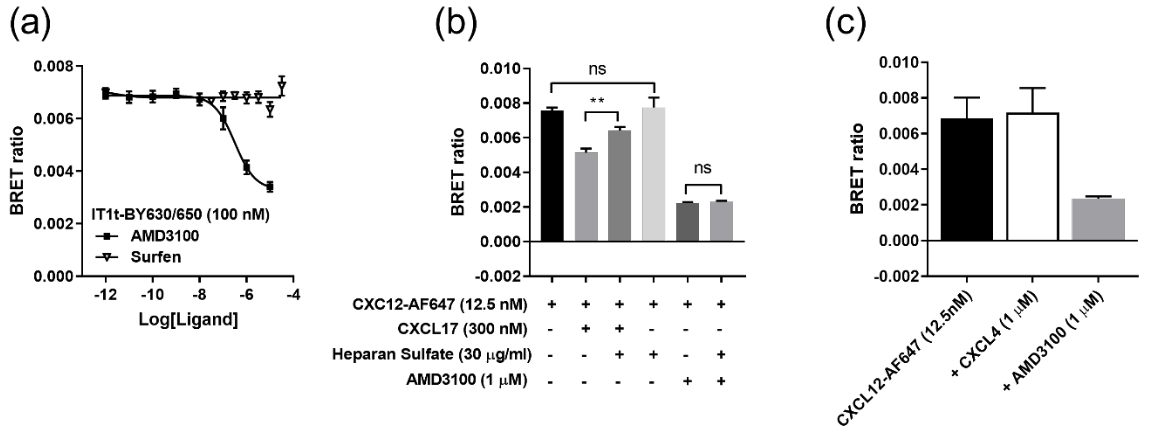


**Supplementary Figure 8: Binders of glycosaminoglycans mimic the effect of CXCL17. (a)** Live HEK293 cells expressing NLuc/CXCR4, were incubated for 1 hr at 37 °C with IT1t-BY630/650 (100 nM) and increasing concentrations of surfen (black triangles, 10 nM – 10 μM) or AMD3100 (black squares, 100 pM – 10 μM). **(b)** Live HEK293 cells expressing genome-edited NLuc/CXCR4 were incubated for 30 min at 37 °C with CXCL12-AF647 (12.5 nM) in the absence of other ligands (black bars), or in the presence of heparan sulfate (30 μg/ml), CXCL17 (300 nM) or AMD3100 (1 μM), or in the presence of CXCL17 (300 nM) or AMD3100 (1 μM) pre-incubated for 1 h with heparan sulfate (30 μg/ml). (**c**) Live HEK293 cells expressing genome-edited NLuc/CXCR4 were incubated for 1 h at 37 °C with CXCL12-AF647 (12.5 nM) in the absence of other ligands (black bars), or in the presence of AMD3100 (1 μM) or CXCL4 (1 μM). Points or bars represent mean ± s.e.m. of four (**a and c**), five or six (**c**) individual experiments performed in duplicate. **, p<0.01 and ns, not statistically significant calculated by one-way ANOVA with Tukey's multiple comparisons test.

**Supplementary Table 1: CXCL17 orthologues used for conservation analysis**

| UniProt ID | Species |
|---|---|
| Q6UXB2 | Homo sapiens (Human) |
| Q5UW37 | Mus musculus (Mouse) |
| H2QGG7 | Pan troglodytes (Chimpanzee) |
| H2NZ12 | Pongo abelii (Sumatran orangutan) (Pongo pygmaeus abelii) |
| D4A875 | Rattus norvegicus (Rat) |
| G1RII2 | Nomascus leucogenys (Northern white-cheeked gibbon) (Hylobates leucogenys) |
| A0A2R8Z5Y3 | Pan paniscus (Pygmy chimpanzee) (Bonobo) |
| A0A337SWE4 | Felis catus (Cat) (Felis silvestris catus) |
| A0A2K6L964 | Rhinopithecus bieti (Black snub-nosed monkey) (Pygathrix bieti) |
| A0A1U7R423 | Mesocricetus auratus (Golden hamster) |
| E2R9V6 | Canis lupus familiaris (Dog) (Canis familiaris) |
| A0A2K6ERT1 | Propithecus coquereli (Coquerel's sifaka) (Propithecus verreauxi coquereli) |
| A0A2K6TQQ2 | Saimiri boliviensis boliviensis (Bolivian squirrel monkey) |
| A0A2K5QSN6 | Cebus capucinus imitator |
| G3QEQ3 | Gorilla gorilla gorilla (Western lowland gorilla) |
| G1LEE3 | Ailuropoda melanoleuca (Giant panda) |
| A0A0D9S176 | Chlorocebus sabaeus (Green monkey) (Cercopithecus sabaeus) |
| A0A0A0MWN4 | Papio anubis (Olive baboon) |
| A0A2K6PII8 | Rhinopithecus roxellana (Golden snub-nosed monkey) (Pygathrix roxellana) |
| A0A2K6CE26 | Macaca nemestrina (Pig-tailed macaque) |
| A0A2K5J518 | Colobus angolensis palliatus (Peters' Angolan colobus) |
| A0A2K5EJ57 | Aotus nancymaae (Ma's night monkey) |
| A0A2K5NDE8 | Cercocebus atys (Sooty mangabey) (Cercocebus torquatus atys) |
| A0A2K5YG25 | Mandrillus leucophaeus (Drill) (Papio leucophaeus) |
| F7DM35 | Callithrix jacchus (White-tufted-ear marmoset) |
| G7PXR6 | Macaca fascicularis (Crab-eating macaque) (Cynomolgus monkey) |
| K7GSG9 | Sus scrofa (Pig) |
| A0A452FVG1 | Capra hircus (Goat) |
| A0A2U3WHI5 | Odobenus rosmarus divergens (Pacific walrus) |
| A0A383YZ12 | Balaenoptera acutorostrata scammoni (North Pacific minke whale) |
| A0A340XH93 | Lipotes vexillifer (Yangtze river dolphin) |
| A0A3Q7MES7 | Callorhinus ursinus (Northern fur seal) |
| A0A2Y9TBG3 | Physeter macrocephalus (Sperm whale) (Physeter catodon) |

| | |
|---|---|
| A0A2Y9HKW8 | Neomonachus schauinslandi (Hawaiian monk seal) (Monachus schauinslandi) |
| A0A341DBJ7 | Neophocaena asiaeorientalis asiaeorientalis (Yangtze finless porpoise) |
| A0A384DLG4 | Ursus maritimus (Polar bear) (Thalarctos maritimus) |
| A0A3Q7VJ80 | Ursus arctos horribilis |
| A0A0P6J2P1 | Heterocephalus glaber (Naked mole rat) |
| A0A1S3ACN0 | Erinaceus europaeus (Western European hedgehog) |
| A0A091EE70 | Fukomys damarensis (Damaraland mole rat) (Cryptomys damarensis) |
| A0A2K5VQK7 | Macaca fascicularis (Crab-eating macaque) (Cynomolgus monkey) |
| A0A2Y9L467 | Enhydra lutris kenyoni |
| A0A485MDS2 | Lynx pardinus (Iberian lynx) (Felis pardina) |
| F7EWH0 | Macaca mulatta (Rhesus macaque) |
| K9IY53 | Desmodus rotundus (Vampire bat) |

Data obtained from Uniprot (Consortium, 2018) and limited to 119 amino acid CXCL17 orthologues.

**Supplementary Table 2: Gains and filter setting used for data collection**

| Figure | Filter | Gain |
|---|---|---|
| Figure 1a | 535-30 / 475-30 | 3600/3500 |
| Figure 1b | 535-30 / 475-30 | 3600/3500 |
| Figure 1c | 535-30 / 475-30 | 3000/2600 |
| Figure 1e | 535-30 / 475-30 | 3000/2600 |
| Figure 1f | 535-30 / 475-30 | 3000/2600 |
| Figure 2a | 535-30 / 475-30 | 3600/3200 |
| Figure 2b | 535-30 / 475-30 | 3600/3200 |
| Figure 2c | 535-30 / 475-30 | 3400/2400 |
| Figure 2d | 535-30 / 475-30 | 3600/3200 |
| Figure 3a | 610-LP / 460-80 | 2600/1900 |
| Figure 3b | 610-LP / 460-80 | 2600/1900 |
| Figure 3d | 610-LP / 460-80 | 3200 / 1800 |
| Figure 3e | 610-LP / 460-80 | 3800 / 2200 |
| Figure 3f | 610-LP / 460-80 | 3600/3200 |
| Figure 3g | 610-LP / 460-80 | 2600/1900 |
| Figure 4 | 535-30 / 475-30 | 3600/3200 |
| Figure 5b | 610-LP / 460-80 | 3600/3200 |
| Figure 5c | 610-LP / 460-80 | 3600/3200 |
| Figure 5d | 610-LP / 460-80 | 3800/2200 |
| Figure 5e | 610-LP / 460-80 | 3600/3200 |
| Figure 6 | 610-LP / 460-80 | 3200 / 1800 |
| Supp. Figure 1a | 535-30 / 475-30 | 3400/2800 |
| Supp. Figure 1b | 535-30 / 475-30 | 3600/3500 |
| Supp. Figure 1c | 535-30 / 475-30 | 3600/3500 |
| Supp. Figure 2 | 535-30 / 475-30 | 3600/3500 |
| Supp. Figure 3 | 535-30 / 475-30 | 3600/3500 |
| Supp. Figure 4a | 610-LP / 460-80 | 3000/2400 |
| Supp. Figure 4b | 610-LP / 460-80 | 3600/3200 |
| Supp. Figure 5a | Total luminescence | 2800 |
| Supp. Figure 5b | 610-LP / 460-80 | 3600/3200 |
| Supp. Figure 6b | 610-LP / 460-80 | 3600/3200 |
| Supp. Figure 7a | 610-LP / 460-80 | 3600/3200 |
| Supp. Figure 7b | 610-LP / 460-80 | 3600/3200 |
| Supp. Figure 7c | 610-LP / 460-80 | 3600/3200 |
